# Intermediately Methylated Regions in Normal Cells Are Epimutation Hotspots in Cancer

**DOI:** 10.1101/2025.09.18.677221

**Authors:** Mohamed Mahgoub, Haley Abel, Nidhi Davarapalli, Heidi Struthers, Emilee Kotnik, Brittany Johnson, Christopher Markovic, Catrina Fronick, Robert Fulton, Michael P. Meers, David H. Spencer

## Abstract

DNA methylation is altered in all cancers, but the mechanisms responsible for these changes are not well understood. Using data from 100 primary samples, we show that regions with intermediate methylation in normal hematopoietic cells are hotspots for stable, clonal epimutations in acute myeloid leukemia. Analysis of hematopoietic stem cell clones demonstrated that intermediate methylation at epimutation hotspots reflects random allele-specific methylation and expression at the single cell level, representing a previously unrecognized form of somatically acquired imprinting. We identified somatically imprinted regions in other tissues and show they are epimutation hotspots in other cancer types. This demonstrates that random allele-specific methylation is both a general property of normal cells, and a vulnerability that renders them susceptible to cooperating events and clonal selection during cancer.

## Main text

Cancer is caused by heritable somatic events that endow neoplastic cells with a selective advantage. While the most extensively studied somatic events in cancer are genetic mutations, cytosine methylation of DNA is also heritable through cell divisions and exhibits tissue-specific changes in malignant vs. normal cells. These include genome-wide methylation patterns present in most cancer types(*1–3*) and signatures that reflect a tumor’s cell of origin, subtype and the effects of somatic mutations(*4–9*). Genome-wide studies have shown that abnormally methylated CpGs in cancer frequently cluster to form differentially methylated regions (DMRs)(*4*, *5*). However, the factors that determine why certain regions acquire these changes are poorly understood. This has made it difficult to define the mechanisms responsible for cancer-associated methylation changes and distinguish early events that could initiate cancer from alterations that are consequences of it(*4*, *10*).

Epimutations are spontaneous, heritable epigenetic abnormalities that can affect gene activity and cause human disease, including cancer(*11*, *12*). The best-characterized epimutations are inherited events that cause the imprinting disorders Beckwith-Wiedemann, Angelman, and Prader-Willi syndromes(*13*, *14*), or predispose patients to cancer via silencing of tumor suppressors(*15*, *16*). Epimutations can also be somatically acquired, most notably in cancer. These can also silence tumor suppressors, producing a second epigenetic ‘hit’ in tumors with a co-occurring somatic mutation(*17–19*). Although the mechanisms and timing of somatic epimutations are not well understood, there is evidence they may pre-exist in normal tissues(*20–24*) and could therefore be selected for during cancer development.

Although methylation in cancer has been extensively studied, most analysis has focused on changes associated with clinical or molecular features(*4*, *6*, *25–29*). Such approaches may be less sensitive to detecting spontaneous epimutations, which could be independent of these variables. Here, we used an unbiased analysis to identify somatically acquired DNA methylation epimutations in acute myeloid leukemia (AML). Surprisingly, these events frequently occurred in regions with intermediate methylation in normal hematopoietic cells, which harbor blocks of CpGs that exhibit random allele-specific methylation in single cells. We demonstrate that methylation at these blocks affects chromatin and gene expression, and that “somatically imprinted” regions are epimutation hotspots in AML and other cancers.

### Intermediately methylated regions in normal hematopoietic cells are hotspots for altered methylation in AML

To identify DNA methylation changes in AML that could be somatically acquired epimutations, we compared whole-genome bisulfite sequencing (WGBS) data from 100 primary AML samples to WGBS from 16 purified hematopoietic cell populations from 7 healthy donors(*4*) (Figure 1A, S1A). Pairwise comparisons were performed between each AML sample to pooled data from stem/progenitor and mature myeloid cells (see Methods) to detect common and rare events and account for maturation-related methylation changes in AML. Between 752 and 19,965 autosomal differentially methylated regions (DMRs) with a methylation difference >0.3 were identified in each sample, for a total of 575,869 DMRs (Figure 1B, S1B). Merging of all DMRs produced 76,752 unique regions, of which 33,181 (43.2%) were “private DMRs” in a single AML sample, and 43,571 (56.8%) regions that overlapped DMRs identified in at least two samples (Figure 1C). Applying a threshold of ≥25 AML samples resulted in 6,076 regions overlapping recurrent DMRs that we defined as DMR “hotspots”, which included regions near genes with important roles in hematopoiesis and AML (e.g., *HOXA3*, Figure 1D).

**Fig. 1.**
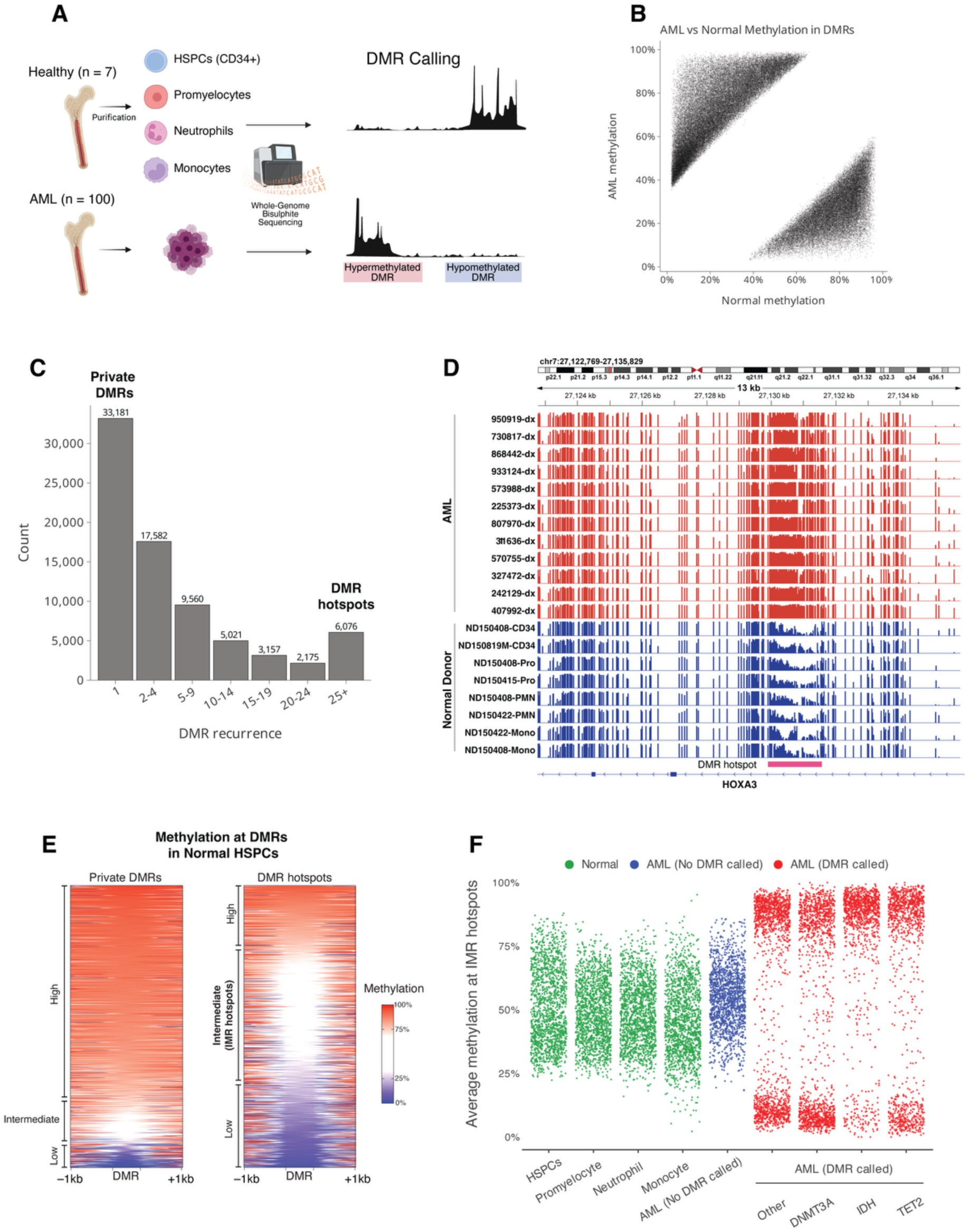
Recurrent differentially methylated regions in AML are enriched for intermediately methylated regions in normal hematopoietic cells. **(A)** Analysis of differentially methylated regions (DMRs) in 100 primary AML samples. WGBS data from each AML was compared to a pooled dataset of 16 hematopoietic cell populations purified from bone marrow aspirates from 7 distinct healthy donors. (**B**) Scatter plot showing the methylation level of each DMR identified in all AML samples versus mean methylation in normal hematopoietic cells. (**C**) Recurrence of DMRs across AMLs after merging overlapping regions from all samples. Private DMRs are regions with differential methylation in one AML sample, while DMR hotspots are regions overlapping highly recurrent DMRs identified in at least 25 AML samples. (**D**) DNA methylation at a DMR hotspot in the *HOXA3* locus from normal hematopoietic cells and selected AML samples. (**E**) Heatmap showing methylation in normal hematopoietic stem/progenitors (HSPCs) at regions overlapping private DMRs identified in only one AML sample (left) and DMR hotspots (25+ samples) (right). DMRs are length-normalized and shown with 1 kb of flanking sequence. (**F**) Methylation levels at intermediately methylated regions in normal hematopoietic cells that are hotspots for differential methylation in AML samples (IMR hotspots). Each point shows the mean methylation of IMR hotspot in the indicated normal hematopoietic cell population (green), AMLs with no DMR called at the hotspot (blue), and AMLs in which a DMR was called (red). AML samples with a DMR at the hotspots are grouped by the presence of mutations in selected genes (minimum of 2 AML samples with the DMR hotspot per mutation group).

We next investigated the normal epigenetic features at hotspots by annotating them with chromatin states(*30*) from hematopoietic stem/progenitor cells (HSPCs) (Figure S1C, see Methods). Hotspots were >10-fold enriched for bivalent chromatin and transcriptional initiation compared to private DMRs, which were evenly distributed across all states (Figure S1D-F). Interestingly, states with the highest enrichment for hotspots had intermediate methylation in normal HSPCs, an uncommon genomic feature that marks tissue-specific regulatory elements(*31*). Direct analysis showed that 48% of hotspots had intermediate methylation in normal HPSCs, and there was a 5-fold enrichment for intermediately methylated CpGs in hotspots compared to their genome-wide frequency (Figure 1E).

We next defined hematopoietic-specific intermediately methylated regions (IMRs) with methylation between 0.3 and 0.7 in normal hematopoietic cells and evaluated the hotspots overlapping these IMRs (i.e., “IMR hotspots”) for features of somatic epimutations. Interestingly, methylation at IMR hotspots remained consistently intermediate throughout myeloid maturation and was present in promyelocytes, monocytes, and neutrophils (Figure 1F), and it was also stable between presentation and relapse in 6 AML patients (Figure S1H). Further, AML samples with DMRs at IMR hotspots showed consistent hyper- or hypomethylation at each hotspot, even in samples with mutations in *DNMT3A*, *IDH1*, *IDH2*, or *TET2* (Figures 1F, S1I) that cause directional methylation changes in AML(*4–6*). This implies that DMRs at IMR hotspots are not simply driven by these mutations. Finally, there was a modest but significant negative correlation between methylation at promoter-associated IMR hotspots and expression of nearby genes, supporting their functional role in gene regulation (Figure S1J). Together this demonstrates that intermediately methylated regions in normal cells are hotspots for methylation changes in AML that resemble somatically acquired epimutations.

### Intermediately methylated regions contain CpG blocks with allelic methylation in AML

Intermediate methylation could be due to cellular heterogeneity, discordant methylation across CpGs, or allele-specific methylation (Figure 2A). To distinguish between these possibilities, we analyzed fragment-level methylation (*32*, *33*) at IMR hotspots in WGBS data from purified normal hematopoietic cells. This showed that methylation at consecutive CpGs was highly correlated at IMR hotspots compared to non-hotspot DMRs with intermediate methylation (p<10^-10^, Figure 2B, S2A). We next tested for allele-specific methylation (ASM) at IMR hotspots with heterozygous SNPs in the normal and AML samples. Surprisingly, ASM was rare in normal samples but more common in AML, especially in samples that retained intermediate methylation (Figure 2C, S2B-C). To further explore this observation, we used Oxford Nanopore sequencing (ONT) to obtain haplotype-resolved methylation from 10 primary AML samples and CD34+ HSPCs or immature (lineage marker-negative) bone marrow cells from four healthy donors (Figure 2D). We identified 1,264-4,497 autosomal regions of ASM per AML sample, after excluding regions in imprinted genes, near structural variants, or with high SNP density; in comparison, ∼300 ASM regions were identified in normal samples (Figure S2D). Interestingly, 30% of the IMR hotspots overlapped an ASM region in the ONT data from AML samples, supporting our findings from WGBS.

**Fig. 2.**
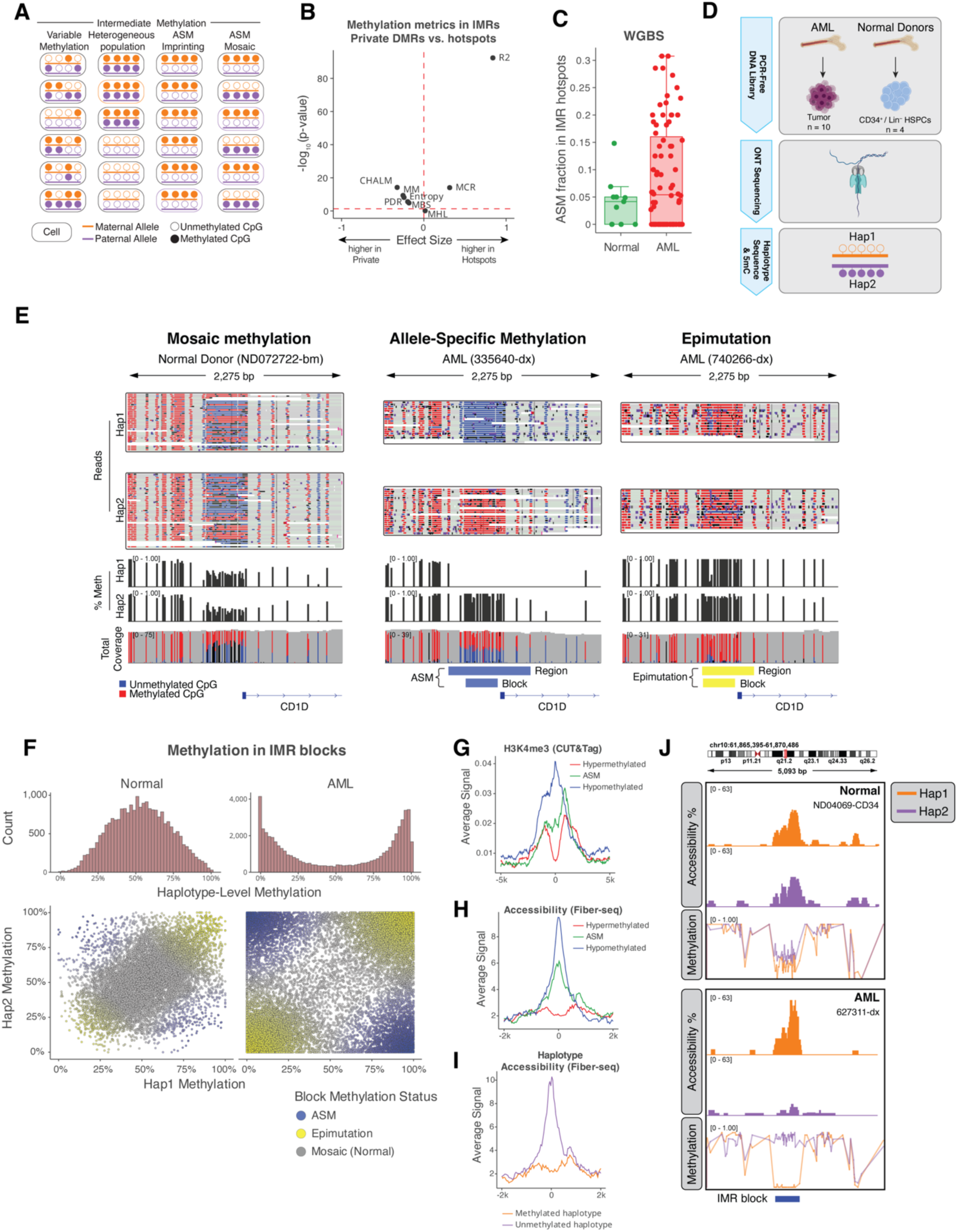
Intermediately methylated regions in normal hematopoietic cells display allelic methylation patterns in AML. **(A)** Models of possible fragment-level methylation patterns at intermediately methylated regions (IMRs). **(B)** Comparison of WGBS fragment-level methylation metrics at IMRs overlapping DMR hotspots vs. non-hotspot DMRs in normal hematopoietic cells. Each point represents a different metric, with the x-axis indicating the effect size (standardized difference in means between hotspots and non-hotspots) and the y-axis indicating statistical significance as –log₁₀(p-value) from a t-test comparing the two groups. Metrics include PDR (Proportion of Discordant Reads), Entropy (Shannon entropy), CHALM (Cell Heterogeneity-Adjusted cLoNal Methylation), R² (linkage disequilibrium R squared), MHL (Methylation Haplotype Load), MCR (Methylation Concordance Ratio), and MBS (Methylation Block Score). **(C)** Fraction of IMR hotspots exhibiting allele-specific methylation (ASM) per sample from WGBS in normal hematopoietic cells and AML samples, calculated among all IMR hotspots with a heterozygous SNP and sufficient coverage. Only samples with at least 5 informative IMR hotspots are included. **(D)** ONT sequencing and haplotype-resolved methylation analysis of normal HSPCs (N=4) and AML samples (N=10). **(E)** Example of haplotype-resolved methylation in an IMR hotspot (at the promoter of *CD1D*), showing mosaic methylation in normal hematopoietic cells, allele-specific methylation (ASM) in AML sample 335640, and full methylation (epimutation) in AML sample 740266. Tracks show the DMR region identified in each AML sample vs. normal cells using DSS R package and the “IMR block” defined by correlation of fragment-level CpG methylation using ONT data(*34*). **(F)** ONT haplotype-level methylation at IMR blocks in all normal HSPCs (N=4) and AML samples (N=10). Top panels show the distribution haplotype-level methylation at IMR blocks. Bottom panels show the average methylation of haplotype 1 versus haplotype 2 for each IMR block. **(G and H)** Aggregated H3K4me3 CUT&Tag signal (G) and chromatin accessibility (Fiber-seq) (H) at IMR blocks in AML samples, with blocks grouped by methylation status (hypomethylated, intermediately methylated, or hypermethylated). **(I)** Chromatin accessibility (Fiber-seq) at IMR blocks with intermediate methylation (green in panel H) and exhibiting ASM in AML samples, comparing accessibility between the methylated and unmethylated haplotypes. **(J)** Example of haplotype-resolved methylation and chromatin accessibility at an IMR block in an intergenic region on chromosome 10 (chr10:61,866,122-61,869,606).

We next used the long-read ONT methylation data from all AML samples and normal HSPCs to identify blocks of consecutive, highly correlated CpGs(*34*) that may be driving allelic methylation at IMR hotspots. We also included published IMRs(*31*) and ASM regions from the ONT data in this analysis, which we reasoned could share similar genomic features. This resulted in 11,534 total methylation haplotype blocks, of which 1,575 had consistently intermediate methylation in normal HSPCs (between 0.3 and 0.7), which we defined as “IMR blocks”. These blocks contained a mean of 12 CpGs, spanned 186 bp, and showed highly correlated—but mosaic—patterns in HSPCs (Figure 2E). Analysis of haplotype-level methylation at all IMR blocks from HSPCs showed that both haplotypes had intermediate methylation, matching the aggregate methylation levels from ONT and WGBS (Figure 2F). In contrast, most IMR blocks in AML samples either had ASM (Figure 2E and 2F: indicated in blue) or were fully methylated/unmethylated on both alleles, reflecting an epimutation (Figure 2E and 2F: indicated in yellow). These findings demonstrate the unique properties of IMR blocks, which have mosaic haplotype-level methylation in normal cells that converts to ASM or fully methylated/unmethylated epimutations in AML.

### Methylation at IMR blocks is associated with differential chromatin activity

We next investigated whether methylation at IMR blocks is associated with differential histone modifications using CUT&Tag and haplotype-level chromatin accessibility via Fiber-seq(*35*). IMR blocks in HSPCs possessed both H3K27 and H3K4 trimethylation, consistent with the bivalent chromatin states at hotspots from the HSPC-defined chromHMM model (Figure S2E). In AML samples, both modifications were negatively correlated with methylation at IMR blocks, with ASM blocks having intermediate signal consistent with allelic histone modification differences (Figure 2G, S2F). Likewise, a negative correlation between methylation and accessibility at IMR blocks was observed via Fiber-seq in two AML samples (Figure 2H). This was also apparent at the haplotype level, with unmethylated haplotypes having higher accessibility in aggregate and at individual regions (Figure 2I, 2J). Overall, these findings are consistent with previous analysis of histone modifications at IMRs(*31*) and support a functional relationship where unmethylated alleles at IMR blocks have greater accessibility and increased histone methylation compared to methylated blocks.

### Allelic methylation at IMR blocks is stable over time and marks clonal hematopoiesis

To determine the stability of allelic methylation at IMR blocks, we used ONT sequencing to analyze paired presentation/relapse samples from 3 AML patients who relapsed after chemotherapy with a related AML clone (Figure 3A, S3A). We compared IMR blocks with allelic methylation (i.e., ASM or a methylated/unmethylated epimutation) between presentation and relapse samples for each patient (using synchronized haplotype designations), which demonstrated that >92.5% of the ASM blocks and >98.9% of the blocks with epimutations were stable at relapse. We next tested whether allelic methylation at IMR blocks is specific to leukemic cells by analyzing remission samples from two of the AML patients. Whole-genome sequencing(*36*) confirmed a molecular remission in patient 895870, while patient 627311 had a persistent *DNMT3A* mutation and several noncoding exome variants, indicating the presence of clonal hematopoiesis (Figure S3B). Patient 895870 had allelic methylation at IMR blocks at presentation that nearly completely normalized in the remission sample before returning at relapse (Figure 3C). Interestingly, allelic methylation only partially normalized in the remission sample from patient 627311 that had clonal hematopoiesis, which instead retained a pattern closer to the presentation and relapse samples (Figure 3D).

**Fig. 3.**
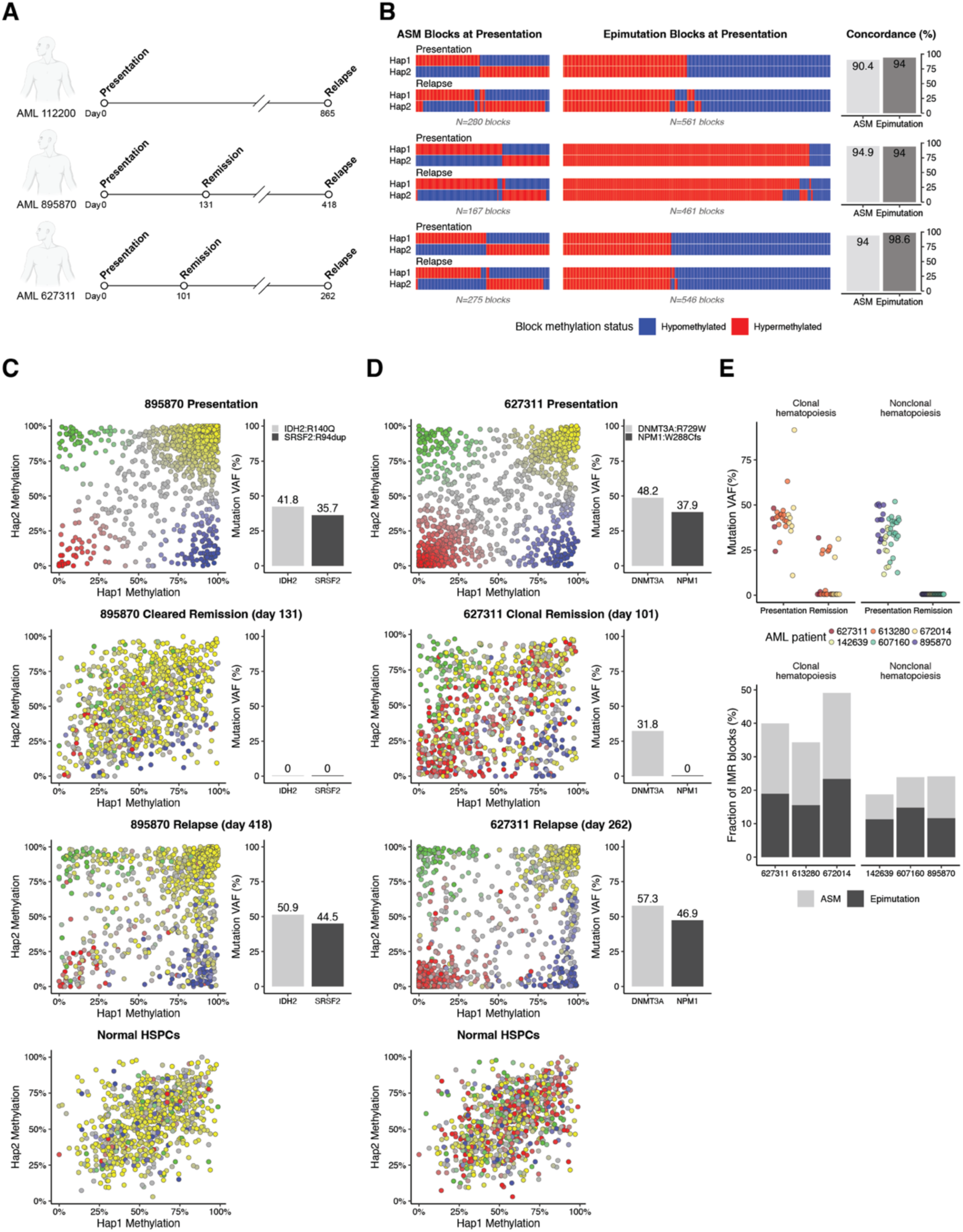
IMR blocks are stable, clonal events in hematopoietic cell populations. **(A)** Longitudinal analysis of methylation at IMR blocks in 3 AML patients (112200, 895870, and 627311) who achieved a complete morphologic remission with standard “7+3” induction chemotherapy. ONT sequencing was performed on the samples at the timepoints indicated by the open circles. **(B)** Heatmap representation of haplotype-level methylation from ONT sequencing of paired presentation/relapse sample pairs. IMR blocks were classified as either having ASM (left) or a fully methylated/unmethylated epimutation (right) via Fisher’s exact test on methylation read counts either between haplotypes (for ASM) or compared to normal HSPCs (for epimutations). Methylated and unmethylated IMR blocks are shown in red and blue, respectively. Only IMR blocks with statistically significant classifications at both presentation and relapse are shown. The percent of concordant ASM and epimutation blocks between presentation and relapse is shown on the right. **(C)** Haplotype-level methylation at IMR blocks from AML patient 895870 at presentation, complete morphologic and molecular remission (i.e., blasts <1% and no leukemia-associated mutations detected), and relapse. Points show the methylation of IMR blocks in haplotype 2 vs. haplotype 1 and are colored by their methylation status in the presentation sample. Bar plots show the variant allele fraction (VAF) of selected recurrent mutations in each sample. Bottom panel shows haplotype-level methylation at the same IMR blocks from a representative CD34+ HSPC sample and are colored based on the methylation status of the block in the presentation AML sample. **(D)** Haplotype-level methylation in AML patient 627311 at presentation, complete morphologic remission (bone marrow blasts <1%), but with a persistent *DNMT3A*, and relapse. Scatter plots are shown as in (C). Bar plots show mutation VAFs in each sample and indicate the presence of a *DNMT3A*^R729W^ variant in the remission sample, consistent with persistent clonal hematopoiesis. **(E)** Correlation between clonal hematopoiesis in AML remission and the presence of ASM and epimutations at IMR blocks from ONT sequencing. Top panels show mutation VAFs at presentation and complete remission for 6 patients, 3 with clonal hematopoiesis in remission (left) and 3 with nonclonal hematopoiesis. Bottom panel shows the fraction of all IMR blocks (N=1,575) that met criteria for being ASM or fully methylated/unmethylated epimutations compared to normal HSPCs in each sample using a Fisher’s exact test on haplotype-level methylation read counts. All remission samples were bone marrow aspirates with normal trilineage hematopoiesis and <1% blasts. See reference (*36*) .

To further examine the relationship between allelic methylation at IMR blocks and clonal hematopoiesis, we used ONT sequencing to measure methylation in additional remission samples from 4 patients who had either complete clearance of somatic mutations or multiple persistent mutations by exome sequencing, indicating clonal hematopoiesis(*36*) (Figure 3E, S3C). Quantification of IMR blocks with allelic methylation showed that patients with clonal hematopoiesis had a significantly higher fraction of allelic blocks compared to samples with complete mutation clearance (41% vs. 22%; p=0.034, Student’s *t*), supporting an association of ASM and epimutations with clonal hematopoiesis.

### Random allele-specific methylation at IMR blocks is somatically acquired in normal hematopoietic stem cells

We reasoned that intermediate methylation at IMR blocks could reflect random ASM that is developmentally established in HSPCs at the single cell level and thus can be observed in clonal AML samples and clonal hematopoiesis but not in bulk cell populations. To test this, we examined IMR block methylation in ONT data from human embryonic stem cells. This showed that most blocks were fully methylated or unmethylated on both haplotypes (Figure S4A), implying that intermediate methylation emerges during hematopoietic development. IMR blocks also reverted to being fully methylated or unmethylated in induced pluripotent stem cells derived from CD34+ HSPCs(*37*) (Figure S4B), indicating these states are intrinsic to pluripotent stem cell identity.

To determine whether mosaic methylation at IMR blocks reflects random ASM, we sorted CD34+/CD38-/CD45RA-HSCs from primary cord blood and expanded bulk HSCs and 11 single-cell HSC clones for analysis via ONT sequencing (Figure 4A). This showed that methylation at most IMR blocks with intermediate methylation in bulk HSCs (684 of 1,575 blocks) did not have mosaic methylation at the haplotype level in single HSC clones: 69% of blocks had ASM in at least one clone and most clones had either ASM or were fully methylated/unmethylated on both haplotypes (Figure 4B). Further, 83% of IMR blocks with ASM in ≥4 clones (N=164) showed ‘switching’ of the methylated haplotype, implying stochastic selection of the methylated state (Figure 4C, 4D). In comparison, haplotype-level methylation was consistent in 98% of regions with ASM in bulk HSCs (Figure 4C), indicating that the random methylation is unlikely to be due to technical or biologic variability. IMR blocks with hyper or hypomethylated epimutations in the HSC clones were also random, but to a lesser extent, with 44% of blocks switching between full hyper and hypomethylated epimutations (Figure S4C, S4D). Most of the epimutation blocks had increased methylation, an effect previously observed in cells forced to rapidly divide in culture(*4*, *38*).

**Fig. 4.**
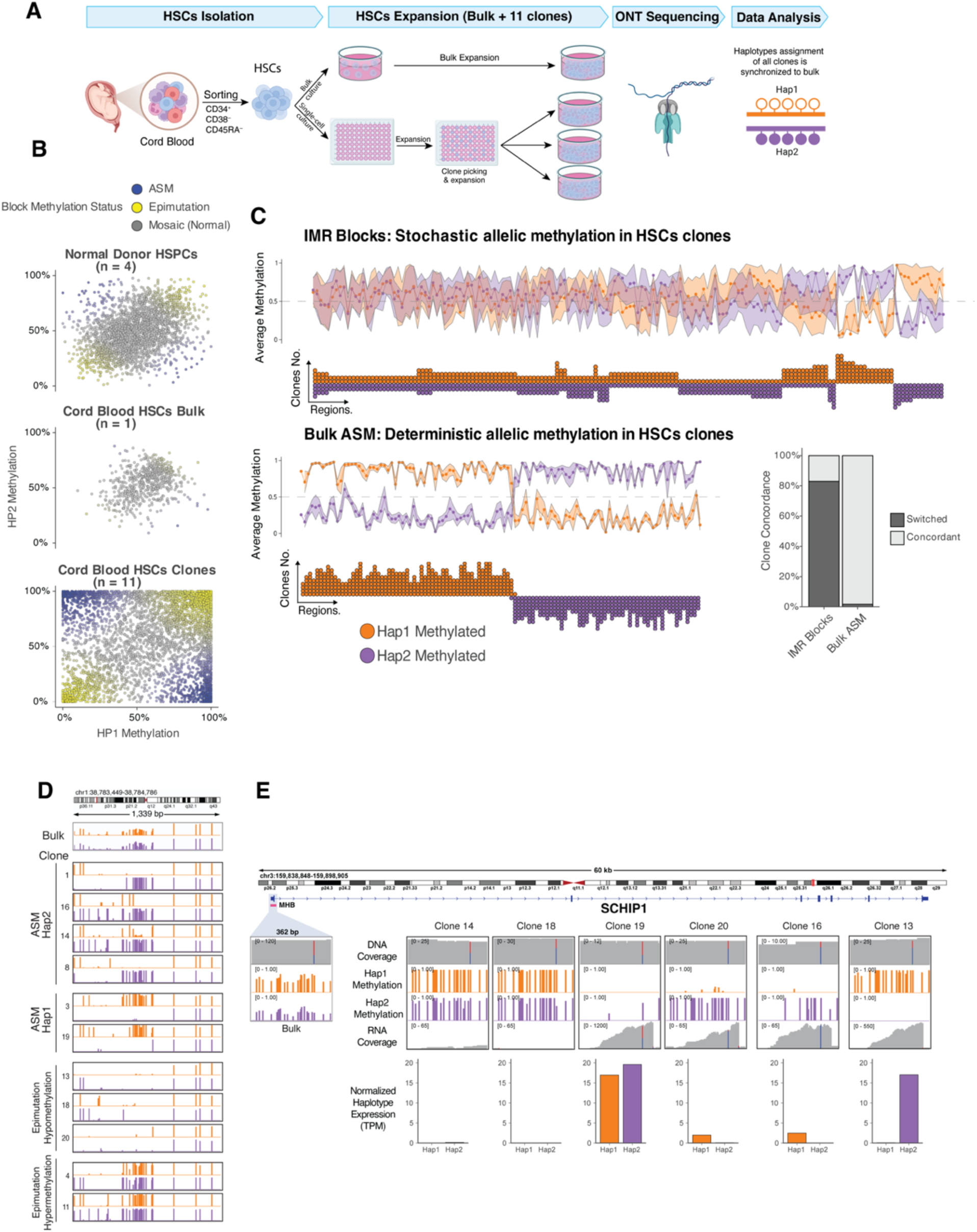
Random allele-specific methylation at IMR blocks is established in single hematopoietic stem cells and is associated with allele-specific gene expression. **(A)** HSC expansion and sequencing to analyze haplotype-level methylation in single-cell-derived HSC clones. **(B)** Haplotype-level methylation at IMR blocks in normal donor HSPCs, bulk cord blood HSCs, and single-cell-derived HSC clones. Each point corresponds to the average methylation of one IMR block within a single clone. Only IMR blocks with intermediate methylation (0.3–0.7) in the bulk HSCs were included. **(C)** Haplotype-level methylation patterns in single-cell-derived HSC clones at IMR blocks exhibiting intermediate methylation in bulk HSCs (top, labelled “IMR Blocks”) and regions showing allele-specific methylation (ASM) in bulk HSCs (bottom left, “Bulk ASM”). In both panels, line plots display the average methylation of haplotype 1 (orange) and haplotype 2 (purple) across all clones for each region. The dot plots below indicate the number of individual clones in which Haplotype 1 or Haplotype 2 was methylated, which is indicated by the Y-axis and the dot color. The bar plot (bottom right) quantifies the percentage of regions where the methylated allele is concordant across clones versus switched (i.e., random) for both IMR blocks and bulk ASM regions. Only regions with ASM observed in at least four clones were included in this analysis. **(D)** Example of an IMR block exhibiting random methylation patterns in HSC clones. The tracks display haplotype-level methylation at the block that shows intermediate methylation in bulk HSCs. Individual clones demonstrate random allele-specific methylation (ASM) on either haplotype 1 (orange) or haplotype 2 (purple), or an epimutation showing either fully hypomethylated or hypermethylated states. **(E)** Allele-specific methylation is associated with allele-specific expression in HSC clones at an IMR block located within an internal promoter of the *SCHIP1* gene. For several representative clones, tracks display DNA coverage, haplotype 1 methylation (orange), haplotype 2 methylation (purple), and RNA coverage. The bar plots quantify normalized, haplotype-specific expression in Transcripts Per Million (TPM). RNA Reads are assigned to haplotypes based on a heterozygous SNP, which is indicated by the vertical blue/grey (haplotype 1) and red (haplotype 2) lines in the coverage tracks. While DNA coverage is consistently biallelic, the level of RNA expression from each haplotype is inversely correlated with its methylation status, demonstrating that higher methylation is linked to transcriptional repression.

We next tested whether random ASM could be linked to allele-specific expression (ASE) by performing bulk RNAseq on 6 HSC clones. Although we were limited to examining only genes with coding SNPs near IMR blocks, we identified a block in an internal promoter of *SCHIP1* with intermediate methylation in bulk HSCs that was either fully methylated/unmethylated or had ASM in all 6 clones (Figure 4E). Haplotype-level expression of *SCHIP1* isoforms using this promoter was variable and correlated with IMR block haplotype methylation. Similar effects were seen for *GATA2*, a critical gene in hematopoietic development previously shown to have ASE in AML(*17*, *22*); several ASM regions were present at the *GATA2* locus in our data, but these did not meet all IMR block criteria. However, *GATA2* had striking monoallelic expression in two clones, and the expression level correlated with haplotype-level methylation in a block-like region (Figure S4E), indicating that both ASM and ASE at *GATA2* are pre-existing states in normal HSCs. Together these observations imply that IMR blocks like those at the *SCHIP1* and *GATA2* loci undergo a form of somatically acquired “imprinting” during hematopoietic development.

### Intermediately methylated blocks in multiple normal tissues are epimutation hotspots in cancer

We were next interested in whether IMR blocks in other tissues displayed epigenetic features like those in hematopoietic cells and were associated with abnormal methylation in tumors. For this analysis, we used data from a recent study of methylation in normal tissues that defined tissue-specific CpG blocks with bimodal methylation (i.e., IMR blocks) (*39*, *40*), as well as separate studies of primary tumors using WGBS or ONT(*41–44*) (Figure 5A). Interestingly, the published set of IMR blocks were enriched for bivalent chromatin and transcriptional start sites in all tissues, similar to the blocks we identified in hematopoietic cells (Figure 5B, see also Figure S1). Likewise, analysis of haplotype-level methylation from primary tumors showed that these blocks lost intermediate methylation and frequently had ASM. Haplotype-level analysis showed that 25-71% of IMR blocks with retained intermediate methylation had ASM across tumor types, further indicating that ASM in these regions is a property of clonal cell populations (Figure 5C, 5D).

**Fig. 5.**
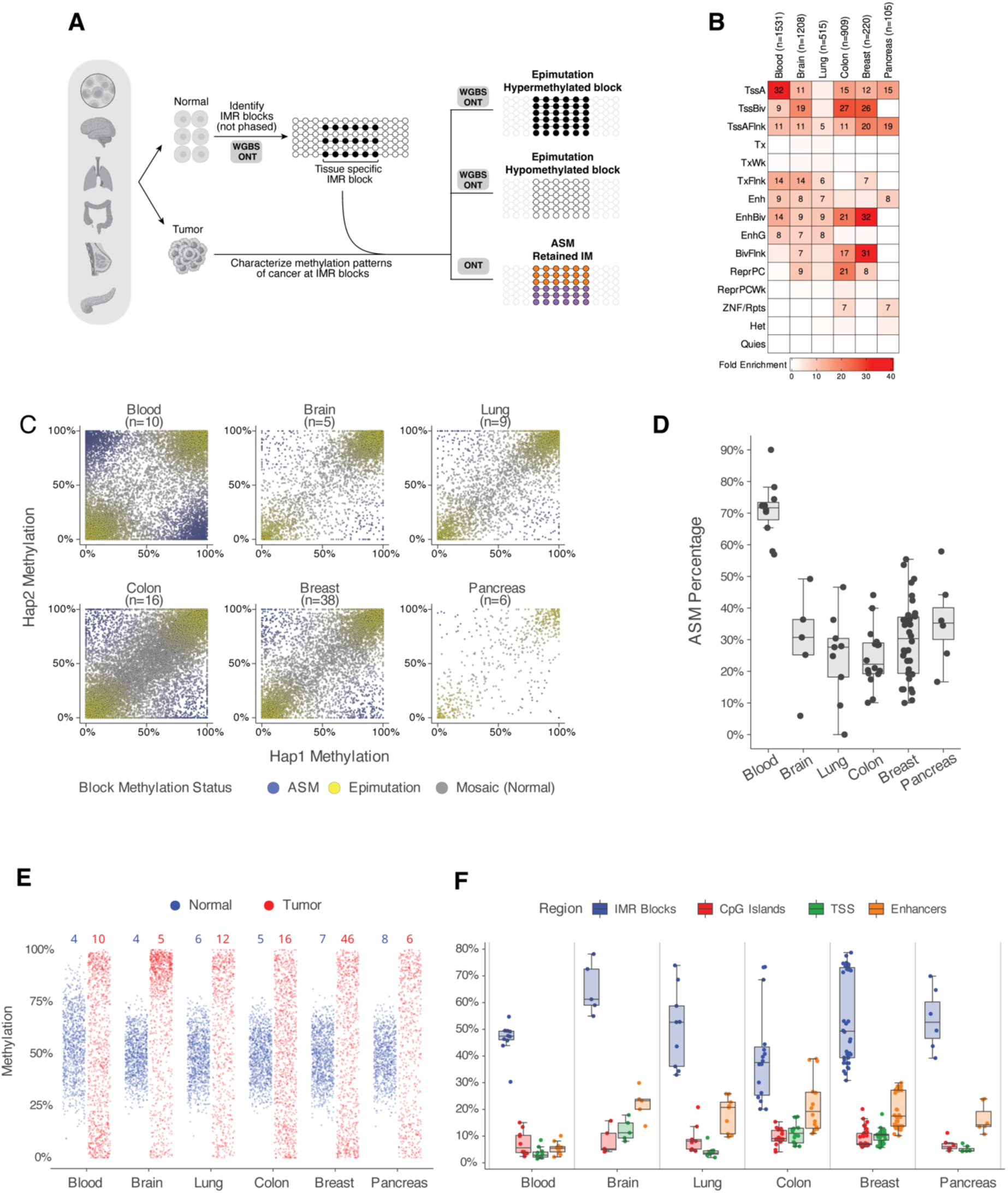
Tissue-specific IMRs in normal cells are hotspots for epimutations in cancer. **(A)** Identifying methylation changes at tissue-specific intermediately methylated regions (IMRs) in multiple cancers. IMRs for hematopoietic cells were identified using ONT data from this study, while IMRs for other tissues were defined using a normal tissue WGBS atlas(*39*). For each tissue, epimutations (hyper- or hypo-methylation) were identified by comparing tumor samples to their normal counterparts. Allele-specific methylation (ASM) was called using haplotype-phased ONT data. This analysis integrated data for AML generated in this study with published datasets for other cancers, including ONT(*41*) and WGBS(*42*, *44*) data. For details, see the Methods section. Heatmap showing the fold enrichment of tissue-specific IMRs within tissue-specific chromatin states. For each tissue (columns), the enrichment of its specific IMRs (n = number of blocks) was calculated for each chromatin state (rows) relative to a genomic background. Values displayed represent the fold enrichment for regions with an enrichment of ≥5. Chromatin state annotations were obtained from the Roadmap Epigenomics Consortium(*45*) . **(C)** Comparison of Haplotype 1 versus Haplotype 2 methylation at tissue-specific IMRs in cancer samples using ONT data**. (D)** Percentage of tissue-specific IMRs with retained intermediate methylation (0.3-0.7) that exhibit allele-specific methylation (ASM) per sample in various cancer types, using phased ONT data. Each point represents an individual tumor sample. The box plots summarize the distribution of ASM percentages for each cancer type shown. **(E)** Distribution of average methylation levels at tissue-specific IMRs in normal tissues (blue) and their corresponding primary tumors (red) from ONT data. The numbers at the top of the plot indicate the number of samples included in each group. Each point represents the average methylation of a single IMR block within a sample, with data from multiple samples pooled for each tissue type. For visualization, 1,000 points were randomly sampled for each group. Data for tumor samples and normal blood were derived from ONT, while data for other normal tissues were from WGBS. **(F)** Percentage of various genomic features exhibiting epimutations across different cancer types from ONT data for individual primary tumor samples. Epimutations were defined as a significant difference in aggregate methylation via a Fisher’s exact test on methylation counts for each region between tumor and its respective normal. Boxplots show the percentage of tissue-specific IMRs (IMR Blocks), CpG islands, tissue-specific transcriptional start sites (TSS), and tissue-specific enhancers that have significant count-based differential methylation in each tumor sample compared to its corresponding normal tissue.

Finally, we sought to determine whether IMR blocks are hotspots for altered methylation in solid tumors and could therefore represent epimutations. Indeed, methylation at tissue-specific IMR blocks was frequently altered in both ONT and WGBS tumor data (Figure 5E, S5A). We then quantified the frequency of altered methylation at IMR blocks by comparing block methylation in each tumor sample to its respective normal tissue. For comparison, the same analysis was performed using other genomic regions commonly associated with differential methylation in cancer, including CpG islands, gene promoters and tissue-specific enhancers defined via chromatin states(*30*). Surprisingly, the fraction of IMR blocks with differential methylation was the highest among these regions, with a mean of 40% to 65% having statistically significant differential methylation vs. normal tissue (Figure 5F, S5B). By comparison, 3.6% to 22% of the other regions had differential methylation across tumor types. This demonstrates that tissue-specific CpG blocks with intermediate methylation are “hotspots” for altered methylation in multiple tumor types.

## Discussion

The factors that predispose certain regions to DNA methylation changes in cancer are not well understood. Here we showed that discrete blocks of CpGs with intermediate methylation in normal cells are *a priori* hotspots for methylation changes in AML and multiple solid tumors. In AML, methylation changes at intermediately methylated regions (IMRs) resemble epimutations: they are stable, clonal events that are less dependent on mutation-driven phenotypes and can be observed in non-leukemic cells. Further, intermediate methylation at these blocks in normal hematopoietic cells is caused by random allele-specific methylation at the single cell level. The highly tissue-specific and stable nature of methylation at these regions implies they normally undergo a type of developmental “somatic imprinting”, and epimutations in AML are due to the absence of this allele-specific state. Further, these regions possess active and bivalent chromatin features and occur near important tissue-specific genes, which suggests they are involved in gene regulation and could be subject to selection during AML pathogenesis.

The concept of allele-specific somatic imprinting is consistent with previous epigenetic studies in primary human tissues. In hematopoietic cells, clonal cell populations exhibit heritable, random monoallelic gene expression(*21*, *22*), and separate studies have used methylation signatures as clonal markers(*46*, *47*). The blocks of random allele-specific methylation described here provide a direct link between these observations by identifying discrete loci where methylation can both contribute to gene regulation and encode stable markers of clonal identity. While it is possible that the mosaic methylation pattern of these regions means they are less functionally relevant, their relationship to chromatin activity and expression suggests otherwise. The striking consistency and stability of IMRs in HSCs also implies that establishing intermediate methylation via somatic imprinting could be a defining feature of HSC identity; epimutations could then be caused by a failure or spontaneous loss of imprinting. While the forces that make ASM advantageous are unknown, it could enable precise gene dosage or simultaneous transcription of two closely spaced genes from opposite alleles. Notably, the *GATA2*, *SCHIP1* and *HOXA* loci with ASM all encode multiple overlapping transcripts and could be subject to such constraints.

One consequence of somatic imprinting is that it generates a substrate for selection during cancer development. For example, genes with ASM and monoallelic expression would be vulnerable to somatic mutations as a second hit(*17*, *18*). Likewise, the presence of epimutations is a source of epigenetic heterogeneity that could confer an advantage to some cells. We found that IMR blocks were differentially methylated more often than CpG islands, promoters, and enhancers in multiple cancers, and in AML, individual IMR blocks tended to become either fully methylated or unmethylated, hinting that certain epimutations may be under selection. The discrete nature of these events means they may be more amenable to direct measurements and functional characterization to understand how pre-existing epigenetic variability contributes to the somatic processes that lead to cancer.

## Materials and Methods

### Primary AML and normal hematopoietic cell samples

Primary AML and normal hematopoietic cell samples used in this study were collected from consented subjects using IRB-approved protocols (WashU HRPO#201011766, FHCC #985.03). AML samples were either peripheral blood or bone marrow aspirate specimens obtained from patients with active disease at presentation or relapse and were processed without fractionation, purification, or enrichment via buffy coat preparation into cryopreserved cell suspensions or directly used for DNA extraction (see Tables S1 and S4). Remission samples were unfractionated whole bone marrow aspirates or peripheral blood samples obtained from patients in morphologic complete remission at the timepoints indicated in Figure 3. Normal adult CD34+ HSPCs, promyelocytes, neutrophils, and monocytes from healthy donor bone marrow aspirate samples as described previously(*4*, *5*). Additional HSPC data were generated from two sources. First, we used peripheral blood mobilized CD34+ HSPCs from 3 donors from the Fred Hutchinson Cancer Center (FHCC) Cooperative Centers of Excellence in Hematology Cell Procurement & Processing Core. Second, we enriched for lineage-negative cells from bone marrow aspirates from two healthy adult donors via bead-based depletion of lineage-positive cells (Miltenyi cat# 130-092-211) on an autoMACS instrument (Miltenyi). Finally, commercially available purified CD34+ HSPCs from cord blood samples (STEMCELL) were used for the clonal expansion experiment.

### Cord blood HSC sorting and expansion

Cryopreserved umbilical cord blood CD34+ cells (10^6^ cells from a single donor purchased from STEMCELL Technologies) were thawed in a ThawSTAR instrument (BioLife) and then cultured overnight in 10 wells of a 96-well U-bottom plate in StemSpan media with CD34+ expansion supplement, UM729 (STEMCELL) and 1% pen/strep antibiotic. After overnight culture, cells were collected and stained with CD34-APC (BD Pharmingen, #555824), CD38-PE (BD Pharmingen, #560981), CD45RA-FITC (BD Pharmingen, #561882) and Near-IR LIVE/DEAD stain (Invitrogen) for FACS on a MoFlo instrument. Single CD34+/CD38-/CD45RA-cells were sorted into two 96-well flat-bottom plates seeded with 25,000 cells/well of irradiated (2000 cGy) AFT024 stroma (ATCC) and containing 100 uL of prepared StemSpan media. An additional 30,000 cells with the same immunophenotype were sorted into an additional well for expansion of a bulk (nonclonal) cell population. On days 4 and 10 after sorting 50 uL of prepared StemSpan media was added to all wells. The bulk HSCs were harvested in day 15 of culture and yielded approximately 10^7^ cells. Expanded single-cell clones were transferred to 24-well plates without AFT024 stroma on days 21 and 23 of culture and then expanded in 0.5 mL of prepared StemSpan media for an additional 4 days before harvesting for DNA and RNA extraction.

### Human embryonic stem cell and iPS cell culture

Human WA0, WA09/H9 ESCs and BM9 iPSCs were cultured in mTESR media (STEMCELL) and passaged using dispase before harvesting for DNA extraction.

### Whole-genome bisulfite sequencing data production

WGBS data from 51 AML samples and 15 normal hematopoietic cell populations were repurposed from prior studies(*4*, *5*). Additional datasets were generated as described previously(*5*) using the Zymo Gold Bisulfite conversion (Zymo Research) and Accel DNA methylation kit (IDT) kits to generate WGBS libraries. QC was performed on Agilent Bioanalyzer or Tapestation instruments prior to sequencing on NovaSeq 6000 or X plus instruments to generate 2x150 paired end reads to a target genome coverage of 10x. All WGBS data were processed using the DRAGEN software suite (v4.2.4) to obtain genome-wide 5mC read counts and methylation fractions.

### ONT sequence data production and processing

All DNA for ONT sequencing was extracted using either the NEB Monarch or QIAmp DNA mini kit. DNA was fragmented and size-selected to eliminate small (<5 kbp) fragments and then prepared for sequencing via the LSK114 kit and sequenced using R10.4.1 flowcells on a PromethION sequencer (Oxford Nanopore Technologies, Oxford, UK) with up to two wash/reload cycles. For data obtained from expanded HSCs, adaptive sampling was performed for all IMRs and ASM regions identified in AML samples. Data processing and analysis, including basecalling, was performed via a Nextflow workflow (https://github.com/dhslab/nf-core-wgsnano, commit 8fe4655) using dorado (v0.5.2+7969fab) for basecalling and read mapping, and the PEPPER-Margin-Deepvariant pipeline for phased germline variant calling of SNV and small indels (v0.8.0)(*54*). Structural variants were identified after data processing using Sniffles(*55*). Haplotagging was performed using Whatshap (v2.2)(*56*), which used a common phased VCF file for presentation/remission/relapse samples from AML patients and the bulk HSC data for the HSC expansion experiment to synchronize the haplotype designations.

Mosdepth (v.0.3.3)(*57*) was used to calculate coverage. Combined and haplotype-specific 5mC counts were generated with modkit (https://github.com/nanoporetech/modkit).

### Fiber-seq on normal HSPCs and primary AML samples

Fiber-seq was performed on cryopreserved mobilized CD34+ HSPCs from two donors (obtained from FHCC) and presentation/relapse sample pairs from two AML patients using the published protocol(*41*) with slight modifications. Cells were thawed in a ThawSTAR instrument and 2x10^6^ cells were collected at 250xg for 5 minutes, washed once in 1mL cold PBS and resuspended in 120 uL CUT&TAG nuclear extraction buffer (20 mM HEPES-KOH, pH 7.9, 10 mM KCl, 0.1% Triton X-100, 20% glycerol, and 0.5mM spermidine) and incubated for 10 minutes on ice.

Nuclei were then collected at 350xg for 5 minutes and resuspended in 60 uL Buffer A (15mM Tris-HCl, 15 mM NaCl, 60 mM KCl, and 1 mM EDTA, pH 8.0) and transferred to a PCR tube before adding 1.5 uL 32 mM s-adenosyl methionine (SAM) and 0.5 uL Hia5 enzyme (200U/uL, a gift from A. Stergachis). Reactions were incubated at 25C in a thermocycler for 10 min and then stopped with 6uL of 10% SDS (final concentration of 1%). The lysate was transferred to a 1.5mL tube with wide bore tips and stored at -80C prior to DNA extraction and sequencing. DNA was extracted with a MagAttract DNA extraction kit, fragmented with a megaruptor and size-selected on a HTpippen to 18-22kbp prior to SMRTbell library preparation. Libraries were sequenced on a PacBio Revio instrument to generate CCS reads containing 5mC and kinetic data for prediction of 6mA via Fibertools(*58*). The FIRE pipeline was then used to obtain haplotype-level accessibility for all samples.

### Histone CUT&Tag on CD34+ HSPCs and AML samples

Histone profiling used a CUT&Tag-direct protocol(*59*) with modifications relating to cryo-preserved primary AML cells and the release of tagmented DNA from Concanavalin A coated (Con-A) magnetic beads. Cryopreserved HSPC and AML cells were thawed at 37°C, washed once in cold PBS, then pelleted for 3 minutes at 600g and resuspended in hypotonic NE1 buffer

(20 mM HEPES-KOH pH 7.9, 10 mM KCl, 0.5 mM spermidine, 10% Triton X-100 and 20% glycerol) followed by incubation on ice for between 45 seconds and 2 minutes for nuclei extraction. The mixture was pelleted for 4 minutes at 1,300g, resuspended in 1× PBS and nuclei were counted using the Trypan blue staining or a ViCell Automated Cell Counter (Beckman Coulter). CUT&Tag-Direct was then carried out as previously described (https://doi.org/10.17504/protocols.io.bcuhiwt6). In brief, nuclei were bound to washed paramagnetic concanavalin A (ConA) beads (Bangs Laboratories) at a concentration of 5 µL washed beads per 50K nuclei and incubated with primary antibody (H3K27me3: Cell Signaling 9733S, H3K4me3: Millipore CS200580) at 4°C overnight in Wash Buffer (10 mM HEPES pH 7.5, 150 mM NaCl, 0.5 mM spermidine and Roche Complete Protease Inhibitor Cocktail) with 2 mM EDTA. Bound nuclei were washed and incubated with 1:100 secondary antibody for 1 hour at room temperature and then washed 2x and incubated in Wash-300 Buffer (Wash Buffer with 300 mM NaCl) with 1:50 5 µM loaded pA–Tn5 for 1 hour at room temperature. Nuclei were washed 2x and tagmented in Wash-300 Buffer with 10 mM MgCl2 for 1 hour at 37 °C and then resuspended sequentially in 50 µl of 10 mM TAPS and 5 µl of 10 mM TAPS with 0.1% SDS and incubated for 1 hour at 58 °C. The resulting suspension was mixed well with 16 µl of 0.9375% Triton X-100, and then 2 µL each of F and R primers and 25 µL 2× NEBNext Master Mix (New England Biolabs) were added for direct amplification with the following conditions: (1) 58 °C for 5 minutes, (2) 72 °C for 5 minutes, (3) 98 °C for 30 seconds, (4) 98 °C 15 seconds, (5) 60 °C for 15 seconds, (6) repeat steps 4–5 13 times, (7) 72 °C for 1 minute and (8) hold at 8 °C. DNA from amplified product was purified via AMPure bead cleanup and resuspended in 25 µl of 10 mM Tris-HCl with 1 mM EDTA. Libraries were assessed using the Qubit (Thermo Fisher Scientific) and TapeStation (Agilent) systems and sequencing was performed on an Illumina NovaSeq Xplus to obtain a minimum of 2 million 2x150 bp paired end reads. All data were analyzed via the nf-core(*60*) CUT&Run analysis pipeline to obtain normalized signal in bigwig format.

### Bulk RNAseq on AML samples and HSC clones

RNA was extracted from 10^6^ cells using QIAquick RNA micro kit (Zymo Research). RNA integrity was assessed by RIN scores by Agilent TapeStation 4150 (Agilent Technologies). Quantification of RNA eluates obtained by Qubit RNA BR Assay kit (Invitrogen-Thermo-Fisher). RNAseq libraries were generated via rRNA/Globin HMR depletion followed by ligation-based library preparation (Watchmaker Genomics). Libraries were assessed using Agilent TapeStation 4150 with High Sensitivity D1000 screen tapes (Agilent Technologies) and Qubit dsDNA HS Assays (Invitrogen-Thermo-Fisher). Sequencing was performed on Illumina NovaSeq Xplus to obtain a minimum of 50 million paired end 2x150 bp reads. Data processing used the Illumina DRAGEN RNA pipeline to obtain normalized gene and transcript-level expression values in transcripts per million (TPM).

### Differentially methylated region (DMR) identification from WGBS data

The DSS(*61*) and bsseq(*62*) packages in R (version 4.3) were used to identify DMRs between each AML sample (N=100) and 17 normal hematopoietic cell methylation datasets (9 CD34+ HSPC samples, 3 promyelocyte samples, 3 neutrophil samples, and 2 monocyte samples, all purified as described above from normal bone marrow aspirates from adults aged 20-50). An FDR cutoff of 0.05 and minimum methylation difference of 0.3 were used within DSS without additional filtering. Analysis of DMR recurrence used *bedtools*(*63*) to merge all DMRs and count overlaps with the DMR set from each AML sample; initial merging and overlapping operations used default settings with no minimum overlap. DMR hotspots were then defined using *bedtools multiintersect* to obtain intervals that had >=25 overlapping DMRs across the 100 AML samples and had at least 5 CpG. Coverage-weighted mean methylation of all ‘multi-intersect’ DMRs was calculated for each AML sample and the pooled normal hematopoietic cell data using *methfast*, a custom c program for aggregating CpG methylation counts over bed intervals (code available upon request). These unmanipulated methylation fractions were used for all figures, statistical calculations, and comparisons, except for the heatmaps in Figure 1E where smoothing was performed to impute missing data for visualization (smoothed values were not used for sorting or determining methylation levels).

### DNA fragment-level and allele-specific methylation analysis from WGBS data

Fragment-level methylation metrics were calculated in private and recurrent DMRs using all normal hematopoietic cell datasets using mHapSuite(*38*). Allele-specific methylation (ASM) analysis was performed by first identifying common single-nucleotide polymorphisms (SNPs) from the 1000 Genomes Project (1000G_phase1.snps.high_confidence.hg38.vcf.gz) that overlapped with IMR hotspots. Sequencing aligned reads covering these “hotspot SNPs” were then partitioned into two groups: those matching the reference allele and those matching the specific alternate allele defined in the 1000 Genomes Project. For the subsequent analysis of an IMR hotspot in an individual sample, each allele group was required to have a minimum of 8 reads, with each read spanning at least 4 CpGs. Individual reads were classified as methylated (≥70% of CpGs methylated) or unmethylated (≤30% of CpGs methylated); reads falling between these thresholds were designated "undetermined". Finally, Fisher’s exact test was used to compare the number of methylated and unmethylated reads between the reference and alternate allele groups at each IMR hotspot. P-values were adjusted for multiple testing using the Benjamini-Hochberg method, and hotspots with an FDR < 0.05 were considered significant ASM sites. The code used for this analysis is available at https://github.com/dhslab/methtools.

### ASM analysis from ONT data

ASM regions were identified from ONT data by first importing haplotype-resolved 5mC counts into *bsseq* objects in R and then calling DMRs between haplotypes with *DSS* using a minimum FDR of 0.05 and methylation difference of 0.4. ASM regions were then further filtered to exclude intervals meeting any of the following criteria: 1) on the X or Y chromosome, 2) overlaps a known imprinted region (N=782, from (*64*, *65*)), 3) occurs within 1,000 bp of an identified structural variant in the ONT data, or 4) overlaps a 10,000 bp region with a SNP density that is in the top 1% across the genome.

### Identification and analysis of IMR blocks in ONT data from normal HSPCs and AML samples

To define IMR blocks we pooled three sets of genomic coordinates: DMR hotspots from our AML WGBS data, ASM regions from our AML ONT data, and known IMRs in normal tissues(*37*). Coordinates from these sources were merged to create a unified set of intervals. Within these intervals, we identified methylation haplotype blocks (MHBs) by adapting a previously described method(*40*) for long read ONT data. Specifically, per-read CpG methylation calls were extracted from the ONT BAM files of four normal donors and 10 AML samples using modkit (https://github.com/nanoporetech/modkit). From this read-level methylation data, we calculated the methylation linkage disequilibrium (mLD) between adjacent CpG sites, measured as the squared Pearson correlation coefficient (*r^2^*), requiring a minimum of 10 reads covering a CpG pair for the calculation. An MHB was then defined as a contiguous region containing at least four consecutive CpG sites where the *r^2^* value between any two adjacent CpGs was ≥ 0.5. The code used for this analysis is available at https://github.com/dhslab/methtools. To identify allele-specific methylation (ASM) and epimutations within IMR blocks using ONT data, we applied Fisher’s exact test. For the ASM analysis, we compared the counts of methylated and unmethylated CpGs between the two haplotypes. For the epimutation analysis, we compared the aggregated counts of methylated and unmethylated CpGs (regardless of haplotype) between AML or HSC samples (for data from expanded HSC bulk cells and clones) and pooled HSPCs from normal donors (n=4). For a block to be included, we required a minimum of four covered CpGs. Furthermore, we mandated a minimum adjusted CpG coverage (defined as total CpG counts divided by the number of CpGs in the block) of 5 per haplotype for ASM and 5 per sample for epimutation. P-values for both analyses were adjusted for multiple testing using the Benjamini-Hochberg method across all evaluable blocks. A significant ASM call was defined by an FDR < 0.05 and an absolute methylation difference between haplotypes > 0.4. A significant epimutation was defined by an FDR < 0.05 and an absolute methylation difference between the test and normal samples > 0.3.

### Histone analysis at IMR blocks

Histone analysis at IMR blocks was performed using H3K4me3 and H3K27me3 CUT&Tag data from 7 AML samples and 3 mobilized CD34+ HSPC samples. All analysis used normalized signal from bigwig files produced by the nf-core pipeline. Mean methylation at each block was classified as being intermediate (i.e., having ASM) or being hyper or hypomethylated on both haplotypes. Aggregate normalized signal for each modification in 200 equally spaced bins was then calculated for a 5 kbp window centered on the block location for analysis by methylation-defined groups.

### Fiber-seq FIRE analysis

Haplotype-resolved chromatin accessibility was assessed using the Fiber-seq Inferred Regulatory Elements (FIRE) analysis pipeline(*66*), available at https://github.com/fiberseq/FIRE. The workflow was run using aligned BAM files as input, and accessibility was quantified as the percentage of accessible fibers per region. Analysis of FIRE signal at IMR blocks used either combined or haplotype-level estimates of percent accessibility and were performed at IMR blocks grouped by methylation status (see histone analysis above).

### Short read genome and exome sequencing of AML samples

Whole-genome sequencing (WGS) was performed on representation and relapse AML samples from patients 112200, 627311, and 895870 with matched normal tissue from skin biopsies. All sequencing was performed using 500ng of DNA input to generate libraries with the Illumina DNA library preparation kit, which were quantified by qPCR and sequenced to obtain a target genome coverage of 60x in 2x150 bp paired end reads on an Illumina NovaSeq Xplus sequencer. Data were analyzed using paired somatic mode with the Illumina DRAGEN software. Identified somatic variants were filtered to retain single nucleotide variants with PASS filter flag and total coverage between 20x and 90x to minimize sequencing artifacts. Whole-exome data for the remission samples was described previously(*42*); variants and their associated variant allele fractions were used directly from that publication.

### IMR block identification in WGBS from normal tissues

To identify IMR blocks in normal tissues (brain, lung, colon, breast, and pancreas), we began with a set of previously defined regions of bimodal methylation “bimodal blocks” from WGBS data(*44*). We then applied a multi-step filtering process to refine these regions and isolate those specific to a single tissue type. First, to ensure consistency within a given tissue, we only retained bimodal blocks that were present in all normal samples for each tissue type. Next, these consistent blocks from all tissue types were merged to create a non-redundant block list (minimum block size of 50 bp; minimum distance between blocks of 50 bp). To isolate truly tissue-specific blocks and exclude germline imprinted regions, this non-redundant list was subjected to a second round of filtering. For each block, we calculated the average methylation within its matched-tissue samples and the average methylation across all other (non-matched) normal samples. A block was kept only if it exhibited intermediate methylation specifically in its matched tissue, defined by two criteria: 1) an average methylation level between 0.3 and 0.7 with a coefficient of variation (CV) ≤ 0.5 within the matched tissue, and 2) an absolute difference in average methylation of ≥ 0.2 between the matched and non-matched tissues. This process resulted in a final set of tissue-specific IMR blocks in hg19 coordinates. For compatibility with the solid-tumor ONT data, a second version of these block coordinates was generated in hg38 using liftOver.

### Analysis of IMR blocks and other regions in WGBS and ONT data from solid tumors

To analyze methylation patterns in solid tumor WGBS and ONT data, we first defined several sets of genomic regions for analysis. These included the IMR blocks described above, CpG islands downloaded from the UCSC genome browser, and tissue-specific TSS and enhancers extracted from Roadmap Epigenomics Consortium ChromHMM annotations (model identifiers: E035 for blood, E073 for brain, E096 for lung, E075 for colon, E027 for breast, E098 for pancreas)(*50*). For the tumor methylation data, we used datasets from GEO: GSE121721 (WGBS: brain), GSE212391 (WGBS: other solid tumors), and GSE270257 (ONT: all solid tumors). For the matched normal samples, we downloaded beta value files from the DNA methylation atlas(*45*). All datasets were processed into a unified CpG-level bedmethyl format; for the normal samples, this conversion was performed using wgbstools(*67*). We then used methfast (https://github.com/dhslab/methfast) to aggregate CpG methylation counts from these bedmethyl files over all genomic regions of interest. For the blood cell type analyses, we used the same WGBS and ONT data for normal and AML samples as previously described. In Figure 5 and Figure S5, these normal hematopoietic cells and AML tumor samples are represented as the ’blood’ normal and tumor category, respectively. Finally, to identify epimutations (in WGBS and ONT) and ASM (in ONT), we applied the same Fisher’s exact test-based approaches used for the hematopoietic cells, which involved comparing aggregated counts between tumor and normal samples for epimutation, and between the two haplotypes for ASM

---

**Fig. S1.**
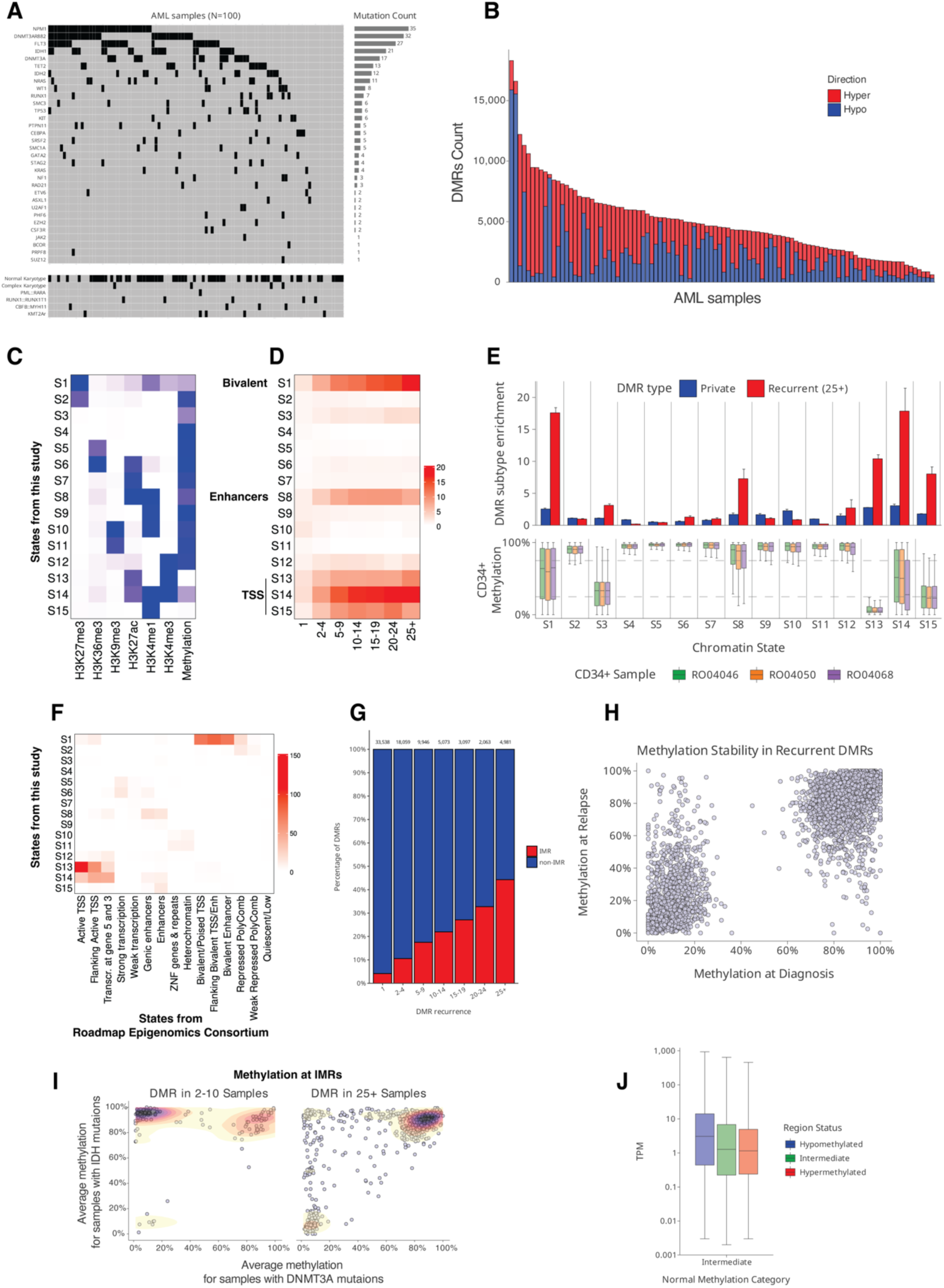
Identification of differentially methylated regions in AML and their enrichment in intermediately methylated regions in normal hematopoietic cells. **(A)** Mutation and karyotype status of AML samples used for WGBS analysis. (**B**) Number of DMRs identified in each AML sample compared to CD34+ cells. **(C)** Emission parameters for states S1 through S15 of a chromHMM model trained on six histone modifications (H3K4me1, H3K4me3, H3K27me3, H3K36me3, H3K9me3, and H3K27ac) and DNA methylation data from purified CD34+ HSPCs. (**D**) Heatmap of DMR enrichment grouped by recurrence frequency in chromatin states defined in HSPCs. (**E**) (Top) Relative enrichment of private and recurrent DMRs and (Bottom) Methylation levels in CD34+ (n=3) at chromatin states. For the methylation analysis, all segments within each state are included. (**F**) Heatmap showing the enrichment mapping of predefined functional genomic regions, annotated in published CD34+ data from the Roadmap Epigenomics Consortium (histone modifications only), within the chromatin states (S1-S15) defined in this study using both histone modifications and DNA methylation in CD34+ cells. (**G**) Overlap percentage of AML DMRs (grouped by recurrence frequency) with previously published intermediately methylated regions (IMRs)(*37*). (**H**) Methylation levels at IMR hotspots with altered methylation (DMRs called), identified in the presentation sample, comparing the methylation levels between presentation and relapse samples from the same patient. (**I**) Comparison of average methylation levels between samples with DNMT3A and IDH mutations, including only those with an identified differentially methylated region (DMR). The analysis distinguishes between two categories of normally intermediately methylated regions (IMRs) based on DMR recurrence: high-recurrence hotspots (DMRs in ≥25 samples) and low-recurrence regions (DMRs in 2–10 samples). Only IMRs with at least two differentially methylated samples in each mutation group were included in the comparison. Regions with low recurrence exhibit methylation patterns that appear to be mutation-driven, while highly recurrent regions are consistently hypermethylated or hypomethylated regardless of mutation type. (**J**) RNA expression levels (TPM) of genes with promoters overlapping recurrent IMR hotspots in AML samples, with samples grouped by promoter methylation status into hypomethylated (<30%), intermediately methylated (30–70%), and hypermethylated (>70%) promoter categories. P < 10^-8^ for a difference in expression across these groups, based on an ANOVA.

**Fig. S2.**
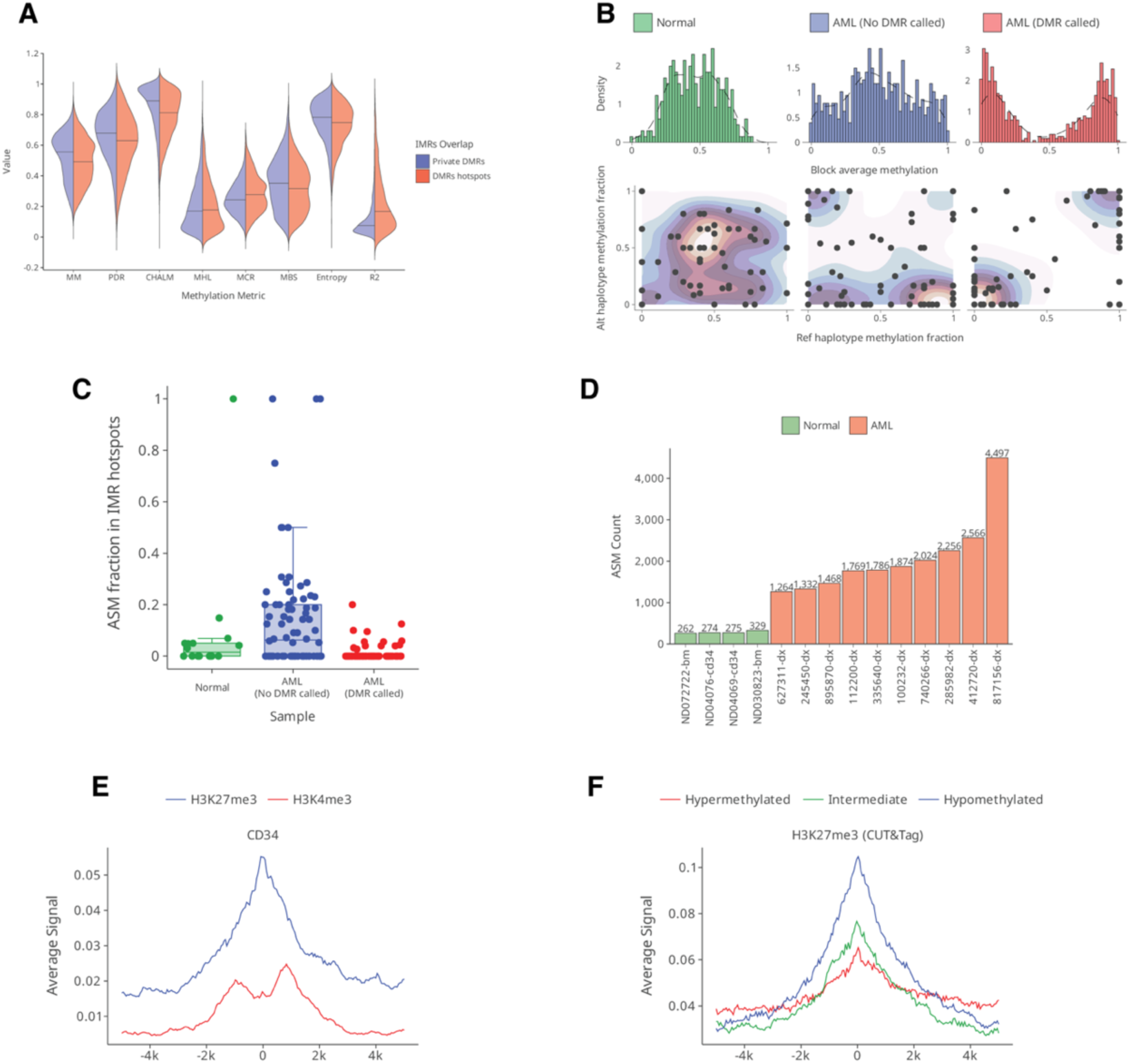
Intermediately methylated regions in normal hematopoietic cells display allelic methylation patterns in AML. **(A)** Fragment-level WGBS methylation metrics statistics at IMRs in normal hematopoietic cells, comparing DMR hotspots vs. non-hotspot DMRs. **(B)** Methylation at IMR hotspots in normal hematopoietic cells, AML samples without differential methylation (no DMR called), and AML samples with DMRs (DMR called). Top: distribution histograms of average methylation at IMR hotspots; bottom: scatter plots comparing the average methylation of reference versus alternate haplotypes for each IMR hotspot with phased WGBS reads above the coverage threshold. **(C)** Fraction of IMR hotspots exhibiting allele-specific methylation (ASM) from WGBS in normal hematopoietic cells, AML without DMRs, and AML with DMRs. **(D)** Number of ASM regions identified from ONT haplotype-resolved methylation using the DSS R package in normal hematopoietic cells and AML samples. **(E)** Aggregated CUT&Tag signal for H3K27me3 and H3K4me3 modifications at IMR blocks in normal hematopoietic cells. **(F)** Aggregated H3K27me3 CUT&Tag signal at IMR blocks in AML samples, with blocks grouped by methylation status (hypomethylated, intermediate, or hypermethylated).

**Fig. S3.**
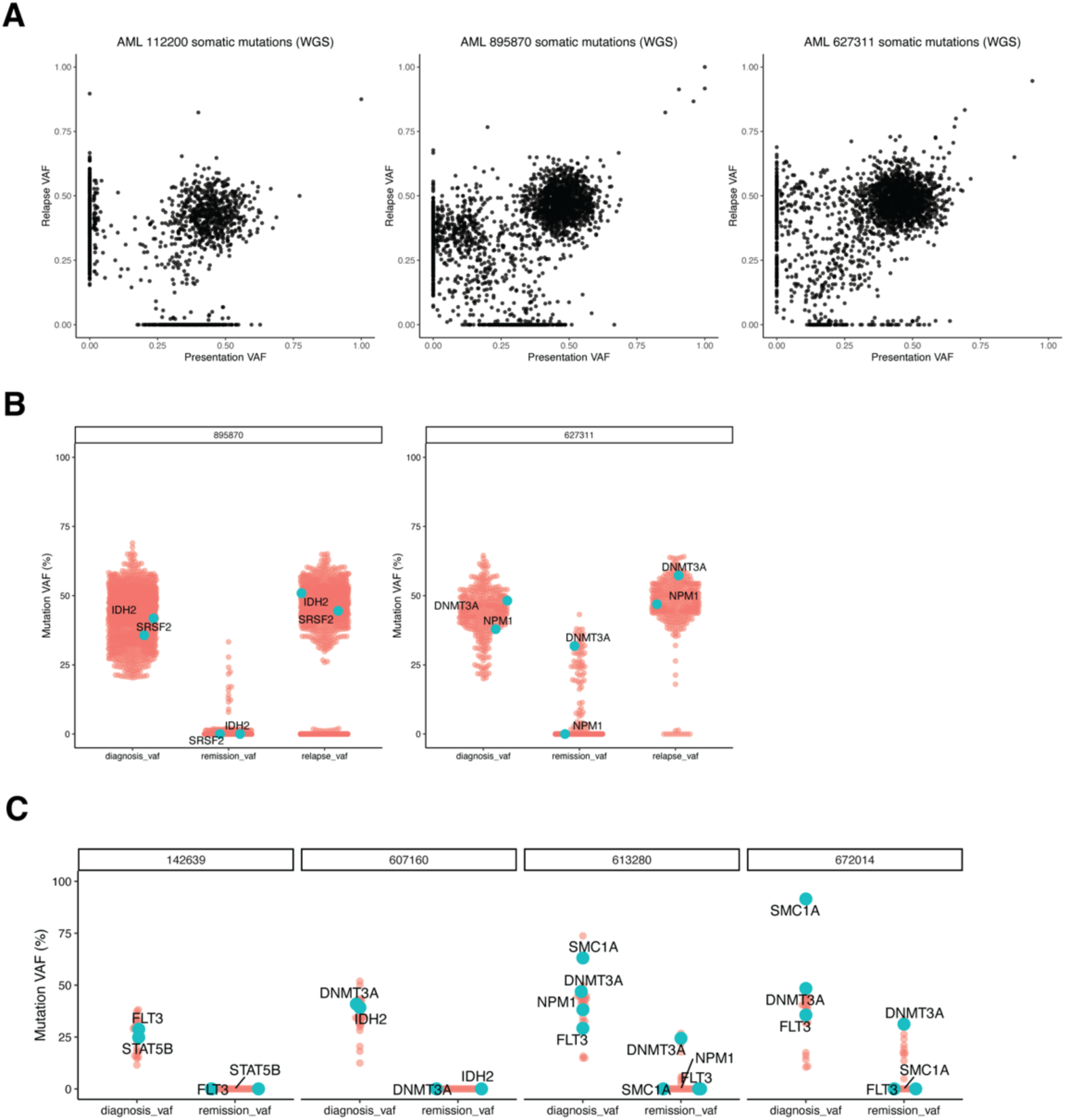
IMR blocks are clonal events in AML and normal hematopoietic cell populations. **(A)** Somatic mutations from whole-genome sequencing of AML patients 112200, 895870, and 627311 at presentation and relapse. Shown are the VAFs at relapse (Y-axis) and presentation (X-axis) demonstrating their clonal relationships. **(B)** VAFs for somatic mutations from AML patient 895870 with nearly complete clearance of all variants in remission (left) vs. AML patient 627311 who had persistent variants, including *DNMT3A*^R729W^, indicating clonal hematopoiesis at the remission timepoint. Both patients had *de novo* AML and were treated with standard “7+3” induction chemotherapy and were in complete morphologic remission at the remission time point. **(C)** Mutation clearance plots from whole-exome sequencing of 4 additional AML patients who achieved a complete morphologic remission after induction chemotherapy(*42*) .

**Fig. S4.**
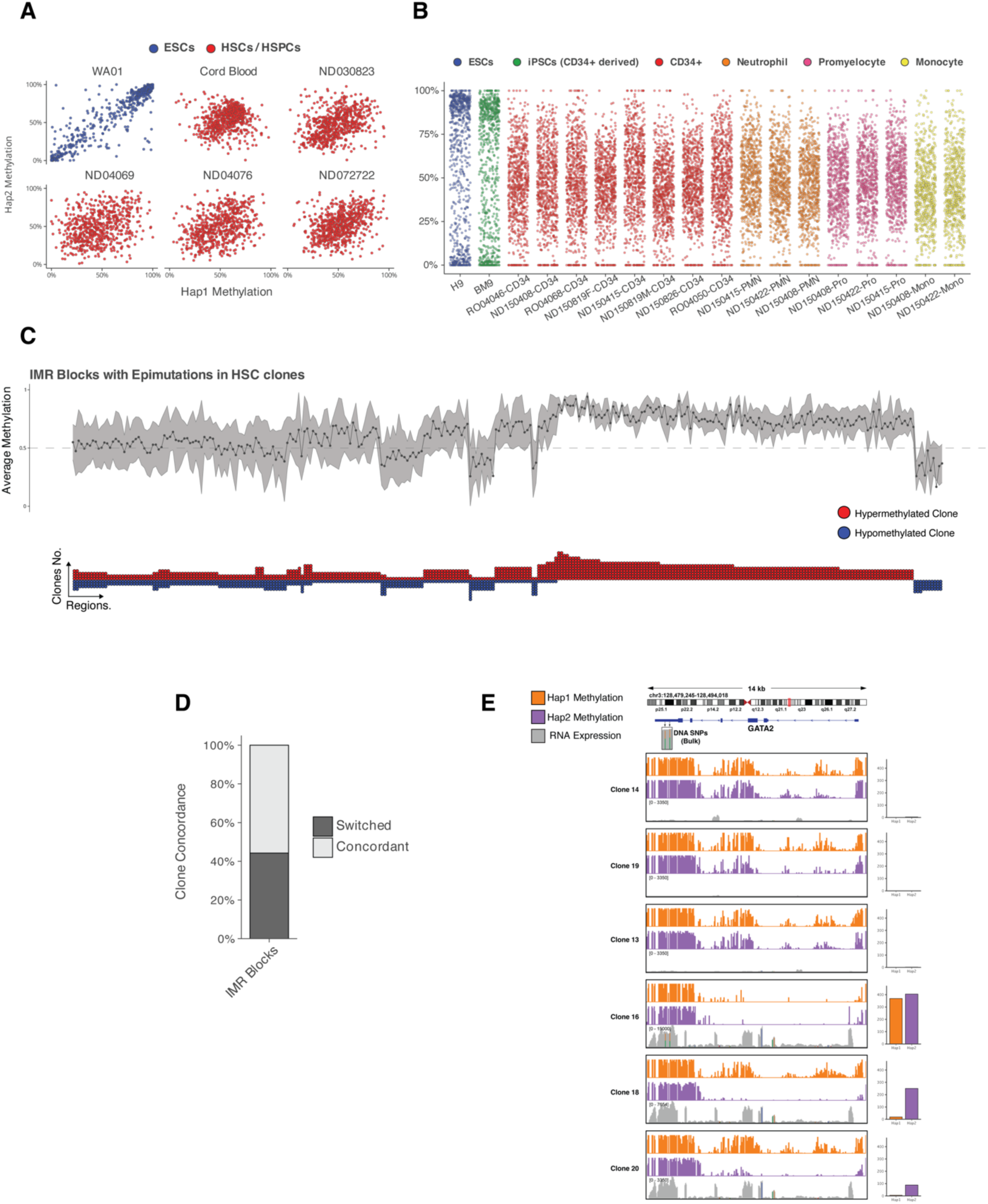
Random allele-specific methylation at IMR blocks is established in single hematopoietic stem cells and is associated with allele-specific gene expression. **(A)** Comparison of haplotype 1 versus haplotype 2 ONT methylation at IMR blocks with intermediate methylation in bulk HSCs. Scatter plots are shown for WA01 embryonic stem cells (ESCs), cord blood HSCs, and normal donor hematopoietic stem and progenitor cells (HSPCs). **(B)** Average WGBS methylation levels at IMR blocks for H9 embryonic stem cells (ESCs), BM9 induced pluripotent stem cells (iPSCs) derived from CD34+ cells, CD34+ cells, and differentiated myeloid lineages. Each point represents the non-phased average WGBS methylation at a single single IMR for the indicated sample. **(C)** Analysis of epimutations in single HSC clones at IMR blocks with intermediate methylation. The top line plot displays the average methylation across all clones for each block. The dot plot below indicates individual clones exhibiting either hyper-methylation (red) or hypo-methylation (blue) epimutations at these blocks relative to normal HSPCs. Only IMR blocks with epimutations observed in at least four clones were included in this analysis. **(D)** Bar plot quantifying the concordance of epimutation type across clones for the IMR blocks shown in (C). “Switched” indicates regions where both hyper- and hypo-methylation events were observed in different clones, while “Concordant” indicates regions where all observed epimutations were of the same type. **(E)** Allele-specific methylation is associated with allele-specific expression at the *GATA2* locus. Tracks for representative single HSC clones show Haplotype 1 methylation (orange), Haplotype 2 methylation (purple), and RNA expression coverage (grey), with two heterozygous SNPs in the last exon used for phasing RNA expression. The bar plots on the right quantify haplotype-specific expression (TPM).

**Fig. S5.**
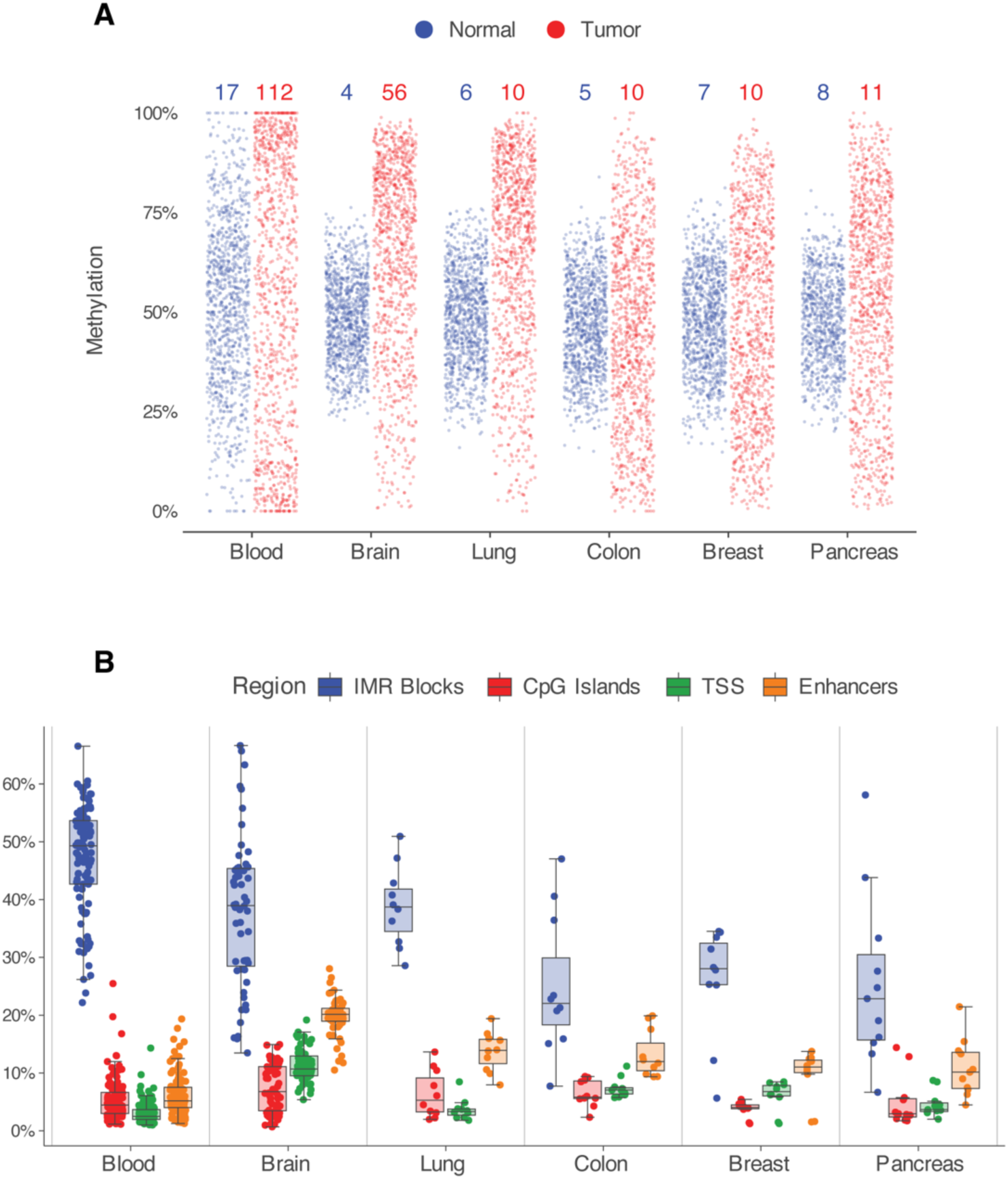
Epimutations at tissue-specific IMRs using WGBS cancer datasets. **(A)** Distribution of average methylation levels at tissue-specific IMRs in normal tissues (blue) and their corresponding primary tumors (red), based on WGBS data. The numbers at the top of the plot indicate the number of samples included in each group. Each point represents the average methylation at a single IMR block within a sample. For visualization, 1,000 points were randomly sampled for each group. **(B)** Percentage of various genomic features exhibiting epimutations across different cancer types, based on WGBS data. The box plots compare the percentage of epimutations within tissue-specific IMRs (IMR Blocks), CpG islands, tissue-specific transcription start sites (TSS), and tissue-specific enhancers. Each point represents an individual tumor sample.

## Acknowledgements

We thank the Miga laboratory at University of California-Santa Cruz for assistance generating the ONT data, M. Jain at Northeastern University for consulting on ONT data analysis, and A. Stergachis at the University of Washington for reagents and assistance with Fiber-seq. Hematopoietic stem cells were purchased from the Cooperative Centers of Excellence in Hematology at the Fred Hutchinson Cancer Center, which is supported in part by NIDDK Grant # DK106829. We are grateful to the patients and research subjects at Alvin J. Siteman Cancer Center at Washington University School of Medicine and Barnes-Jewish Hospital in St. Louis for their participation in the Genomics of AML banking study, which is supported by WashU AML SPORE (P50CA171963, Daniel C. Link, PI), and thank the Siteman Cancer Center for the use of the Siteman Tissue Procurement and Flow Cytometry Cores. The Siteman Cancer Center is supported in part by an NCI Cancer Center Support Grant P30CA091842. We also thank D. Russler-Germain, A. Young, and J. Edwards at Wash U for helpful comments when preparing this manuscript.

## Author contributions

Conceptualization: MM, DHS Resources: DHS

Investigation: MM, HJA, ND, HS, EK, BJ, CM, CF, RF, MPM, DHS

Funding acquisition: DHS Project administration: DHS Visualization: MM, HJA, DHS

Writing – original draft: MM, DHS Writing – review & editing: All authors

## Data and materials availability

Raw and processed genomic data has been deposited to dbGaP (PHS000159) and GEO (accession pending), respectively.

## References

1. A. P. Feinberg, B. Vogelstein, Hypomethylation distinguishes genes of some human cancers from their normal counterparts. Nature 301, 89–92 (1983).

2. A. P. Feinberg, B. Vogelstein, Hypomethylation of ras oncogenes in primary human cancers. Biochem Bioph Res Co 111, 47–54 (1983).

3. M. A. Gama-Sosa, V. A. Slagel, R. W. Trewyn, R. Oxenhandler, K. C. Kuo, C. W. Gehrke, M. Ehrlich, The 5-methylcytosine content of DNA from human tumors. Nucleic Acids Res. 11, 6883–6894 (1983).

4. D. H. Spencer, D. A. Russler-Germain, S. Ketkar, N. M. Helton, T. L. Lamprecht, R. S. Fulton, C. C. Fronick, M. O’Laughlin, S. E. Heath, M. Shinawi, P. Westervelt, J. E. Payton, L. D. Wartman, J. S. Welch, R. K. Wilson, M. J. Walter, D. C. Link, J. F. DiPersio, T. J. Ley, CpG Island Hypermethylation Mediated by DNMT3A Is a Consequence of AML Progression. Cell 168, 801–816.e13 (2017).

5. E. R. Wilson, N. M. Helton, S. E. Heath, R. S. Fulton, J. E. Payton, J. S. Welch, M. J. Walter, P. Westervelt, J. F. DiPersio, D. C. Link, C. A. Miller, T. J. Ley, D. H. Spencer, Focal disruption of DNA methylation dynamics at enhancers in IDH-mutant AML cells. Leukemia 36, 935–945 (2022).

6. J. L. Glass, D. Hassane, B. J. Wouters, H. Kunimoto, R. Avellino, F. E. Garrett-Bakelman, O. A. Guryanova, R. Bowman, S. Redlich, A. M. Intlekofer, C. Meydan, T. Qin, M. Fall, A. Alonso, M. L. Guzman, P. J. M. Valk, C. B. Thompson, R. Levine, O. Elemento, R. Delwel, A. Melnick, M. E. Figueroa, Epigenetic Identity in AML Depends on Disruption of Nonpromoter Regulatory Elements and Is Affected by Antagonistic Effects of Mutations in Epigenetic Modifiers. Cancer Discov 7, 868–883 (2017).

7. D. Capper, D. T. W. Jones, M. Sill, V. Hovestadt, D. Schrimpf, D. Sturm, C. Koelsche, F. Sahm, L. Chavez, D. E. Reuss, A. Kratz, A. K. Wefers, K. Huang, K. W. Pajtler, L. Schweizer, D. Stichel, A. Olar, N. W. Engel, K. Lindenberg, P. N. Harter, A. K. Braczynski, K. H. Plate, H. Dohmen, B. K. Garvalov, R. Coras, A. Hölsken, E. Hewer, M. Bewerunge-Hudler, M. Schick, R. Fischer, R. Beschorner, J. Schittenhelm, O. Staszewski, K. Wani, P. Varlet, M. Pages, P. Temming, D. Lohmann, F. Selt, H. Witt, T. Milde, O. Witt, E. Aronica, F. Giangaspero, E. Rushing, W. Scheurlen, C. Geisenberger, F. J. Rodriguez, A. Becker, M. Preusser, C. Haberler, R. Bjerkvig, J. Cryan, M. Farrell, M. Deckert, J. Hench, S. Frank, J. Serrano, K. Kannan, A. Tsirigos, W. Brück, S. Hofer, S. Brehmer, M. Seiz-Rosenhagen, D. Hänggi, V. Hans, S. Rozsnoki, J. R. Hansford, P. Kohlhof, B. W. Kristensen, M. Lechner, B. Lopes, C. Mawrin, R. Ketter, A. Kulozik, Z. Khatib, F. Heppner, A. Koch, A. Jouvet, C. Keohane, H. Mühleisen, W. Mueller, U. Pohl, M. Prinz, A. Benner, M. Zapatka, N. G. Gottardo, P. H. Driever, C. M. Kramm, H. L. Müller, S. Rutkowski, K. von Hoff, M. C. Frühwald, A. Gnekow, G. Fleischhack, S. Tippelt, G. Calaminus, C.-M. Monoranu, A. Perry, C. Jones, T. S. Jacques, B. Radlwimmer, M. Gessi, T. Pietsch, J. Schramm, G. Schackert, M. Westphal, G. Reifenberger, P. Wesseling, M. Weller, V. P. Collins, I. Blümcke, M. Bendszus, J. Debus, A. Huang, N. Jabado, P. A. Northcott, W. Paulus, A. Gajjar, G. W. Robinson, M. D. Taylor, Z. Jaunmuktane, M. Ryzhova, M. Platten, A. Unterberg, W. Wick, M. A. Karajannis, M. Mittelbronn, T. Acker, C. Hartmann, K. Aldape, U. Schüller, R. Buslei, P. Lichter, M. Kool, C. Herold-Mende, D. W. Ellison, M. Hasselblatt, M. Snuderl, S. Brandner, A. Korshunov, A. von Deimling, S. M. Pfister, DNA methylation-based classification of central nervous system tumours. Nature 555, 469–474 (2018).

8. C. Koelsche, D. Schrimpf, D. Stichel, M. Sill, F. Sahm, D. E. Reuss, M. Blattner, B. Worst, C. E. Heilig, K. Beck, P. Horak, S. Kreutzfeldt, E. Paff, S. Stark, P. Johann, F. Selt, J. Ecker, D. Sturm, K. W. Pajtler, A. Reinhardt, A. K. Wefers, P. Sievers, A. Ebrahimi, A. Suwala, F. Fernández-Klett, B. Casalini, A. Korshunov, V. Hovestadt, F. K. F. Kommoss, M. Kriegsmann, M. Schick, M. Bewerunge-Hudler, T. Milde, O. Witt, A. E. Kulozik, M. Kool, L. Romero-Pérez, T. G. P. Grünewald, T. Kirchner, W. Wick, M. Platten, A. Unterberg, M. Uhl, A. Abdollahi, J. Debus, B. Lehner, C. Thomas, M. Hasselblatt, W. Paulus, C. Hartmann, O. Staszewski, M. Prinz, J. Hench, S. Frank, Y. M. H. Versleijen-Jonkers, M. E. Weidema, T. Mentzel, K. Griewank, E. de Álava, J. D. Martín, M. A. I. Gastearena, K. T.-E. Chang, S. Y. Y. Low, A. Cuevas-Bourdier, M. Mittelbronn, M. Mynarek, S. Rutkowski, U. Schüller, V. F. Mautner, J. Schittenhelm, J. Serrano, M. Snuderl, R. Büttner, T. Klingebiel, R. Buslei, M. Gessler, P. Wesseling, W. N. M. Dinjens, S. Brandner, Z. Jaunmuktane, I. Lyskjær, P. Schirmacher, A. Stenzinger, B. Brors, H. Glimm, C. Heining, O. M. Tirado, M. Sáinz-Jaspeado, J. Mora, J. Alonso, X. G. del Muro, S. Moran, M. Esteller, J. K. Benhamida, M. Ladanyi, E. Wardelmann, C. Antonescu, A. Flanagan, U. Dirksen, P. Hohenberger, D. Baumhoer, W. Hartmann, C. Vokuhl, U. Flucke, I. Petersen, G. Mechtersheimer, D. Capper, D. T. W. Jones, S. Fröhling, S. M. Pfister, A. von Deimling, Sarcoma classification by DNA methylation profiling. Nat. Commun. 12, 498 (2021).

9. B. Giacopelli, M. Wang, A. Cleary, Y.-Z. Wu, A. R. Schultz, M. Schmutz, J. S. Blachly, A.-K. Eisfeld, B. Mundy-Bosse, S. Vosberg, P. A. Greif, R. Claus, L. Bullinger, R. Garzon, K. R. Coombes, C. D. Bloomfield, B. J. Druker, J. W. Tyner, J. C. Byrd, C. C. Oakes, DNA methylation epitypes highlight underlying developmental and disease pathways in acute myeloid leukemia. Genome Res 31, gr.269233.120 (2021).

10. S. Baylin, T. H. Bestor, Altered methylation patterns in cancer cell genomes: Cause or consequence? Cancer Cell 1, 299–305 (2002).

11. M. P. Hitchins, Constitutional epimutation as a mechanism for cancer causality and heritability? Nat. Rev. Cancer 15, 625–634 (2015).

12. B. Horsthemke, Heritable germline epimutations in humans. Nat. Genet. 39, 573–574 (2007).

13. R. L. Poole, L. E. Docherty, A. A. Sayegh, A. Caliebe, C. Turner, E. Baple, E. Wakeling, L. Harrison, A. Lehmann, I. K. Temple, D. J. G. Mackay, O. behalf of the I. C. I. Consortium, Targeted methylation testing of a patient cohort broadens the epigenetic and clinical description of imprinting disorders. Am. J. Méd. Genet. Part A 161, 2174–2182 (2013).

14. K. Buiting, S. Groß, C. Lich, G. Gillessen-Kaesbach, O. El-Maarri, B. Horsthemke, Epimutations in Prader-Willi and Angelman Syndromes: A Molecular Study of 136 Patients with an Imprinting Defect. Am. J. Hum. Genet. 72, 571–577 (2003).

15. M. P. Hitchins, R. L. Ward, Constitutional (germline) MLH1 epimutation as an aetiological mechanism for hereditary non-polyposis colorectal cancer. J. Méd. Genet. 46, 793 (2009).

16. C. M. Suter, D. I. K. Martin, R. L. Ward, Germline epimutation of MLH1 in individuals with multiple cancers. Nat. Genet. 36, 497–501 (2004).

17. R. Mulet-Lazaro, S. van Herk, C. Erpelinck, E. Bindels, M. A. Sanders, C. Vermeulen, I. Renkens, P. Valk, A. M. Melnick, J. de Ridder, M. Rehli, C. Gebhard, R. Delwel, B. J. Wouters, Allele-specific expression of GATA2 due to epigenetic dysregulation in CEBPA double-mutant AML. Blood 138, 160–177 (2021).

18. S. B. Simpkins, T. Bocker, E. M. Swisher, D. G. Mutch, D. J. Gersell, A. J. Kovatich, J. P. Palazzo, R. Fishel, P. J. Goodfellow, MLH1 Promoter Methylation and Gene Silencing is the Primary Cause of Microsatellite Instability in Sporadic Endometrial Cancers. Hum. Mol. Genet. 8, 661–666 (1999).

19. K. O’Neill, E. Pleasance, J. Fan, V. Akbari, G. Chang, K. Dixon, V. Csizmok, S. MacLennan, V. Porter, A. Galbraith, C. J. Grisdale, L. Culibrk, J. H. Dupuis, R. Corbett, J. Hopkins, R. Bowlby, P. Pandoh, D. E. Smailus, D. Cheng, T. Wong, C. Frey, Y. Shen, L. F. Paulin, F. J. Sedlazeck, J. M. T. Nelson, E. Chuah, K. L. Mungall, R. A. Moore, R. Coope, A. J. Mungall, M. K. McConechy, L. M. Williamson, K. A. Schrader, S. Yip, M. A. Marra, J. Laskin, S. J. M. Jones, Long-read sequencing of an advanced cancer cohort resolves rearrangements, unravels haplotypes, and reveals methylation landscapes. medRxiv, 2024.02.20.24302959 (2024).

20. A. Gimelbrant, J. N. Hutchinson, B. R. Thompson, A. Chess, Widespread Monoallelic Expression on Human Autosomes. Science 318, 1136–1140 (2007).

21. O. Stewart, C. Gruber, H. E. Randolph, R. Patel, M. Ramba, E. Calzoni, L. H. Huang, J. Levy, S. Buta, A. Lee, C. Sazeides, Z. Prue, D. P. H. van Konijnenburg, I. K. Chinn, L. A. Pedroza, J. R. Lupski, E. G. Schmitt, M. A. Cooper, A. Puel, X. Peng, S. Boisson-Dupuis, J. Bustamante, S. Okada, M. Martin-Fernandez, J. S. Orange, J.-L. Casanova, J. D. Milner, D. Bogunovic, Monoallelic expression can govern penetrance of inborn errors of immunity. Nature, 1–12 (2025).

22. A. F. A. Seraihi, A. Rio-Machin, K. Tawana, C. Bödör, J. Wang, A. Nagano, J. A. Heward, S. Iqbal, S. Best, N. Lea, D. McLornan, E. J. Kozyra, M. W. Wlodarski, C. M. Niemeyer, H. Scott, C. Hahn, A. Ellison, H. Tummala, S. R. Cardoso, T. Vulliamy, I. Dokal, T. Butler, M. Smith, J. Cavenagh, J. Fitzgibbon, GATA2 monoallelic expression underlies reduced penetrance in inherited GATA2-mutated MDS/AML. Leukemia 32, 2502–2507 (2018).

23. L. Marion-Poll, B. Forêt, D. Zielinski, F. Massip, M. Attia, A. C. Carter, L. Syx, H. Y. Chang, A.-V. Gendrel, E. Heard, Locus specific epigenetic modalities of random allelic expression imbalance. Nat. Commun. 12, 5330 (2021).

24. Q. Deng, D. Ramsköld, B. Reinius, R. Sandberg, Single-Cell RNA-Seq Reveals Dynamic, Random Monoallelic Gene Expression in Mammalian Cells. Science 343, 193–196 (2014).

25. H. Noushmehr, D. J. Weisenberger, K. Diefes, H. S. Phillips, K. Pujara, B. P. Berman, F. Pan, C. E. Pelloski, E. P. Sulman, K. P. Bhat, R. G. W. Verhaak, K. A. Hoadley, D. N. Hayes, C. M. Perou, H. K. Schmidt, L. Ding, R. K. Wilson, D. V. D. Berg, H. Shen, H. Bengtsson, P. Neuvial, L. M. Cope, J. Buckley, J. G. Herman, S. B. Baylin, P. W. Laird, K. Aldape, T. C. G. A. R. Network, Identification of a CpG Island Methylator Phenotype that Defines a Distinct Subgroup of Glioma. Cancer Cell 17, 510–522 (2010).

26. T. J. Ley, C. Miller, L. Ding, B. J. Raphael, A. J. Mungall, A. G. Robertson, K. Hoadley, T. J. Triche, P. W. Laird, J. D. Baty, L. L. Fulton, R. Fulton, S. E. Heath, J. Kalicki-Veizer, C. Kandoth, J. M. Klco, D. C. Koboldt, K.-L. Kanchi, S. Kulkarni, T. L. Lamprecht, D. E. Larson, L. Lin, C. Lu, M. D. McLellan, J. F. McMichael, J. Payton, H. Schmidt, D. H. Spencer, M. H. Tomasson, J. W. Wallis, L. D. Wartman, M. A. Watson, J. Welch, M. C. Wendl, A. Ally, M. Balasundaram, I. Birol, Y. Butterfield, R. Chiu, A. Chu, E. Chuah, H.-J. Chun, R. Corbett, N. Dhalla, R. Guin, A. He, C. Hirst, M. Hirst, R. A. Holt, S. Jones, A. Karsan, D. Lee, H. I. Li, M. A. Marra, M. Mayo, R. A. Moore, K. Mungall, J. Parker, E. Pleasance, P. Plettner, J. Schein, D. Stoll, L. Swanson, A. Tam, N. Thiessen, R. Varhol, N. Wye, Y. Zhao, S. Gabriel, G. Getz, C. Sougnez, L. Zou, M. D. M. Leiserson, F. Vandin, H.-T. Wu, F. Applebaum, S. B. Baylin, R. Akbani, B. M. Broom, K. Chen, T. C. Motter, K. Nguyen, J. N. Weinstein, N. Zhang, M. L. Ferguson, C. Adams, A. Black, J. Bowen, J. Gastier-Foster, T. Grossman, T. Lichtenberg, L. Wise, T. Davidsen, J. A. Demchok, K. R. M. Shaw, M. Sheth, H. J. Sofia, L. Yang, J. R. Downing, G. Eley, Genomic and Epigenomic Landscapes of Adult De Novo Acute Myeloid Leukemia. New Engl J Med 368, 2059–2074 (2013).

27. T. Mazor, A. Pankov, B. E. Johnson, C. Hong, E. G. Hamilton, R. J. A. Bell, I. V. Smirnov, G. F. Reis, J. J. Phillips, M. J. Barnes, A. Idbaih, A. Alentorn, J. J. Kloezeman, M. L. M. Lamfers, A. W. Bollen, B. S. Taylor, A. M. Molinaro, A. B. Olshen, S. M. Chang, J. S. Song, J. F. Costello, DNA Methylation and Somatic Mutations Converge on the Cell Cycle and Define Similar Evolutionary Histories in Brain Tumors. Cancer Cell 28, 307–317 (2015).

28. T. C. G. A. R. Network, Comprehensive, Integrative Genomic Analysis of Diffuse Lower-Grade Gliomas. New Engl J Medicine 372, 2481–2498 (2015).

29. M. E. Figueroa, S. Lugthart, Y. Li, C. Erpelinck-Verschueren, X. Deng, P. J. Christos, E. Schifano, J. Booth, W. van Putten, L. Skrabanek, F. Campagne, M. Mazumdar, J. M. Greally, P. J. M. Valk, B. Löwenberg, R. Delwel, A. Melnick, DNA Methylation Signatures Identify Biologically Distinct Subtypes in Acute Myeloid Leukemia. Cancer Cell 17, 13–27 (2010).

30. J. Ernst, M. Kellis, Chromatin-state discovery and genome annotation with ChromHMM. Nat Protoc 12, 2478–2492 (2017).

31. G. Elliott, C. Hong, X. Xing, X. Zhou, D. Li, C. Coarfa, R. J. A. Bell, C. L. Maire, K. L. Ligon, M. Sigaroudinia, P. Gascard, T. D. Tlsty, R. A. Harris, L. C. Schalkwyk, M. Bilenky, J. Mill, P. J. Farnham, M. Kellis, M. A. Marra, A. Milosavljevic, M. Hirst, G. D. Stormo, T. Wang, J. F. Costello, Intermediate DNA methylation is a conserved signature of genome regulation. Nat. Commun. 6, 6363 (2015).

32. Y. Ding, K. Cai, L. Liu, Z. Zhang, X. Zheng, J. Shi, mHapTk: a comprehensive toolkit for the analysis of DNA methylation haplotypes. Bioinformatics 38, 5141–5143 (2022).

33. Y. Feng, Z. Zhang, Y. Hong, Y. Ding, L. Liu, S. Gao, H. Fang, J. Shi, A DNA methylation haplotype block landscape in human tissues and preimplantation embryos reveals regulatory elements defined by comethylation patterns. Genome Res. 33, 2041–2052 (2023).

34. S. Guo, D. Diep, N. Plongthongkum, H.-L. Fung, K. Zhang, K. Zhang, Identification of methylation haplotype blocks aids in deconvolution of heterogeneous tissue samples and tumor tissue-of-origin mapping from plasma DNA. Nat. Genet. 49, 635–642 (2017).

35. A. B. Stergachis, B. M. Debo, E. Haugen, L. S. Churchman, J. A. Stamatoyannopoulos, Single-molecule regulatory architectures captured by chromatin fiber sequencing. Science 368, 1449– 1454 (2020).

36. M. J. Slade, R. Ghasemi, M. O’Laughlin, T. Burton, R. S. Fulton, H. J. Abel, E. J. Duncavage, T. J. Ley, M. A. Jacoby, D. H. Spencer, Persistent Molecular Disease in Adult Patients With AML Evaluated With Whole-Exome and Targeted Error-Corrected DNA Sequencing. *JCO Precis*. Oncol. 7, e2200559 (2023).

37. K. Hu, J. Yu, K. Suknuntha, S. Tian, K. Montgomery, K.-D. Choi, R. Stewart, J. A. Thomson, I. I. Slukvin, Efficient generation of transgene-free induced pluripotent stem cells from normal and neoplastic bone marrow and cord blood mononuclear cells. Blood 117, e109–e119 (2011).

38. V. L. Wilson, P. A. Jones, DNA Methylation Decreases in Aging But Not in Immortal Cells. Science 220, 1055–1057 (1983).

39. J. Rosenski, A. Peretz, J. Magenheim, N. Loyfer, R. Shemer, B. Glaser, Y. Dor, T. Kaplan, Atlas of imprinted and allele-specific DNA methylation in the human body. Nat. Commun. 16, 2141 (2025).

40. N. Loyfer, J. Magenheim, A. Peretz, G. Cann, J. Bredno, A. Klochendler, I. Fox-Fisher, S. Shabi-Porat, M. Hecht, T. Pelet, J. Moss, Z. Drawshy, H. Amini, P. Moradi, S. Nagaraju, D. Bauman, D. Shveiky, S. Porat, U. Dior, G. Rivkin, O. Or, N. Hirshoren, E. Carmon, A. Pikarsky, A. Khalaileh, G. Zamir, R. Grinbaum, M. A. Gazala, I. Mizrahi, N. Shussman, A. Korach, O. Wald, U. Izhar, E. Erez, V. Yutkin, Y. Samet, D. R. Golinkin, K. L. Spalding, H. Druid, P. Arner, A. M. J. Shapiro, M. Grompe, A. Aravanis, O. Venn, A. Jamshidi, R. Shemer, Y. Dor, B. Glaser, T. Kaplan, A DNA methylation atlas of normal human cell types. Nature, 1– 10 (2023).

41. K. O’Neill, E. Pleasance, J. Fan, V. Akbari, G. Chang, K. Dixon, V. Csizmok, S. MacLennan, V. Porter, A. Galbraith, C. J. Grisdale, L. Culibrk, J. H. Dupuis, R. Corbett, J. Hopkins, R. Bowlby, P. Pandoh, D. E. Smailus, D. Cheng, T. Wong, C. Frey, Y. Shen, E. Lewis, L. F. Paulin, F. J. Sedlazeck, J. M. T. Nelson, E. Chuah, K. L. Mungall, R. A. Moore, R. Coope, A. J. Mungall, M. K. McConechy, L. M. Williamson, K. A. Schrader, S. Yip, M. A. Marra, J. Laskin, S. J. M. Jones, Long-read sequencing of an advanced cancer cohort resolves rearrangements, unravels haplotypes, and reveals methylation landscapes. Cell Genom. 4, 100674 (2024).

42. Z. Zhang, Y. Hong, S. Zhang, X. Zhu, L. Liu, X. Liao, H. Gu, H. Fang, J. Shi, Toward the DNA methylation haplotype map of 11 common solid cancers. Cell Rep. 44, 116197 (2025).

43. A. L. Costa, D. Doncevic, Y. Wu, L. Yang, K. H. Man, A.-S. Spreng, H. Winter, M. N. C. Fletcher, B. Radlwimmer, C. Herrmann, A new IDH-independent hypermethylation phenotype is associated with astrocyte-like cell state in glioblastoma. Genome Biol. 26, 192 (2025).

44. Y. Wu, M. Fletcher, Z. Gu, Q. Wang, B. Costa, A. Bertoni, K.-H. Man, M. Schlotter, J. Felsberg, J. Mangei, M. Barbus, A.-C. Gaupel, W. Wang, T. Weiss, R. Eils, M. Weller, H. Liu, G. Reifenberger, A. Korshunov, P. Angel, P. Lichter, C. Herrmann, B. Radlwimmer, Glioblastoma epigenome profiling identifies SOX10 as a master regulator of molecular tumour subtype. Nat. Commun. 11, 6434 (2020).

45. A. Kundaje, W. Meuleman, J. Ernst, M. Bilenky, A. Yen, A. Heravi-Moussavi, P. Kheradpour, Z. Zhang, J. Wang, M. J. Ziller, V. Amin, J. W. Whitaker, M. D. Schultz, L. D. Ward, A. Sarkar, G. Quon, R. S. Sandstrom, M. L. Eaton, Y.-C. Wu, A. Pfenning, X. Wang, M. C. Liu, C. Coarfa, R. A. Harris, N. Shoresh, C. B. Epstein, E. Gjoneska, D. Leung, W. Xie, R. D. Hawkins, R. Lister, C. Hong, P. Gascard, A. J. Mungall, R. Moore, E. Chuah, A. Tam, T. K. Canfield, R. S. Hansen, R. Kaul, P. J. Sabo, M. S. Bansal, A. Carles, J. R. Dixon, K.-H. Farh, S. Feizi, R. Karlic, A.-R. Kim, A. Kulkarni, D. Li, R. Lowdon, G. Elliott, T. R. Mercer, S. J. Neph, V. Onuchic, P. Polak, N. Rajagopal, P. Ray, R. C. Sallari, K. T. Siebenthall, N. A. Sinnott-Armstrong, M. Stevens, R. E. Thurman, J. Wu, B. Zhang, X. Zhou, N. Abdennur, M. Adli, M. Akerman, L. Barrera, J. Antosiewicz-Bourget, T. Ballinger, M. J. Barnes, D. Bates, R. J. A. Bell, D. A. Bennett, K. Bianco, C. Bock, P. Boyle, J. Brinchmann, P. Caballero-Campo, R. Camahort, M. J. Carrasco-Alfonso, T. Charnecki, H. Chen, Z. Chen, J. B. Cheng, S. Cho, A. Chu, W.-Y. Chung, C. Cowan, Q. A. Deng, V. Deshpande, M. Diegel, B. Ding, T. Durham, L. Echipare, L. Edsall, D. Flowers, O. Genbacev-Krtolica, C. Gifford, S. Gillespie, E. Giste, I. A. Glass, A. Gnirke, M. Gormley, H. Gu, J. Gu, D. A. Hafler, M. J. Hangauer, M. Hariharan, M. Hatan, E. Haugen, Y. He, S. Heimfeld, S. Herlofsen, Z. Hou, R. Humbert, R. Issner, A. R. Jackson, H. Jia, P. Jiang, A. K. Johnson, T. Kadlecek, B. Kamoh, M. Kapidzic, J. Kent, A. Kim, M. Kleinewietfeld, S. Klugman, J. Krishnan, S. Kuan, T. Kutyavin, A.-Y. Lee, K. Lee, J. Li, N. Li, Y. Li, K. L. Ligon, S. Lin, Y. Lin, J. Liu, Y. Liu, C. J. Luckey, Y. P. Ma, C. Maire, A. Marson, J. S. Mattick, M. Mayo, M. McMaster, H. Metsky, T. Mikkelsen, D. Miller, M. Miri, E. Mukame, R. P. Nagarajan, F. Neri, J. Nery, T. Nguyen, H. O’Geen, S. Paithankar, T. Papayannopoulou, M. Pelizzola, P. Plettner, N. E. Propson, S. Raghuraman, B. J. Raney, A. Raubitschek, A. P. Reynolds, H. Richards, K. Riehle, P. Rinaudo, J. F. Robinson, N. B. Rockweiler, E. Rosen, E. Rynes, J. Schein, R. Sears, T. Sejnowski, A. Shafer, L. Shen, R. Shoemaker, M. Sigaroudinia, I. Slukvin, S. Stehling-Sun, R. Stewart, S. L. Subramanian, K. Suknuntha, S. Swanson, S. Tian, H. Tilden, L. Tsai, M. Urich, I. Vaughn, J. Vierstra, S. Vong, U. Wagner, H. Wang, T. Wang, Y. Wang, A. Weiss, H. Whitton, A. Wildberg, H. Witt, K.-J. Won, M. Xie, X. Xing, I. Xu, Z. Xuan, Z. Ye, C. Yen, P. Yu, X. Zhang, X. Zhang, J. Zhao, Y. Zhou, J. Zhu, Y. Zhu, S. Ziegler, A. E. Beaudet, L. A. Boyer, P. L. D. Jager, P. J. Farnham, S. J. Fisher, D. Haussler, S. J. M. Jones, W. Li, M. A. Marra, M. T. McManus, S. Sunyaev, J. A. Thomson, T. D. Tlsty, L.-H. Tsai, W. Wang, R. A. Waterland, M. Q. Zhang, L. H. Chadwick, B. E. Bernstein, J. F. Costello, J. R. Ecker, M. Hirst, A. Meissner, A. Milosavljevic, B. Ren, J. A. Stamatoyannopoulos, T. Wang, M. Kellis, A. Kundaje, W. Meuleman, J. Ernst, M. Bilenky, A. Yen, A. Heravi-Moussavi, P. Kheradpour, Z. Zhang, J. Wang, M. J. Ziller, V. Amin, J. W. Whitaker, M. D. Schultz, L. D. Ward, A. Sarkar, G. Quon, R. S. Sandstrom, M. L. Eaton, Y.-C. Wu, A. R. Pfenning, X. Wang, M. Claussnitzer, Y. Liu, C. Coarfa, R. A. Harris, N. Shoresh, C. B. Epstein, E. Gjoneska, D. Leung, W. Xie, R. D. Hawkins, R. Lister, C. Hong, P. Gascard, A. J. Mungall, R. Moore, E. Chuah, A. Tam, T. K. Canfield, R. S. Hansen, R. Kaul, P. J. Sabo, M. S. Bansal, A. Carles, J. R. Dixon, K.-H. Farh, S. Feizi, R. Karlic, A.-R. Kim, A. Kulkarni, D. Li, R. Lowdon, G. Elliott, T. R. Mercer, S. J. Neph, V. Onuchic, P. Polak, N. Rajagopal, P. Ray, R. C. Sallari, K. T. Siebenthall, N. A. Sinnott-Armstrong, M. Stevens, R. E. Thurman, J. Wu, B. Zhang, X. Zhou, A. E. Beaudet, L. A. Boyer, P. L. D. Jager, P. J. Farnham, S. J. Fisher, D. Haussler, S. J. M. Jones, W. Li, M. A. Marra, M. T. McManus, S. Sunyaev, J. A. Thomson, T. D. Tlsty, L.-H. Tsai, W. Wang, R. A. Waterland, M. Q. Zhang, L. H. Chadwick, B. E. Bernstein, J. F. Costello, J. R. Ecker, M. Hirst, A. Meissner, A. Milosavljevic, B. Ren, J. A. Stamatoyannopoulos, T. Wang, M. Kellis, Integrative analysis of 111 reference human epigenomes. Nature 518, 317–330 (2015).

45. M. Chen, R. Fu, Y. Chen, L. Li, S.-W. Wang, High-resolution, noninvasive single-cell lineage tracing in mice and humans based on DNA methylation epimutations. Nat. Methods, 1–11 (2025).

46. M. Scherer, I. Singh, M. M. Braun, C. Szu-Tu, P. S. Sanchez, D. Lindenhofer, N. A. Jakobsen, V. Körber, M. Kardorff, L. Nitsch, P. Kautz, J. Rühle, A. Bianchi, L. Cozzuto, R. Frömel, S. Beneyto-Calabuig, C. Lareau, A. T. Satpathy, R. Beekman, L. M. Steinmetz, S. Raffel, L. S. Ludwig, P. Vyas, A. Rodriguez-Fraticelli, L. Velten, Clonal tracing with somatic epimutations reveals dynamics of blood ageing. Nature 643, 478–487 (2025).

